# Chronic cholesterol administration to the brain supports complete and long-lasting cognitive and motor amelioration in Huntington’s disease

**DOI:** 10.1101/2022.08.26.505426

**Authors:** Giulia Birolini, Marta Valenza, Ilaria Ottonelli, Francesca Talpo, Lucia Minoli, Andrea Cappelleri, Mauro Bombaci, Claudio Caccia, Caterina Canevari, Arianna Trucco, Valerio Leoni, Alice Passoni, Monica Favagrossa, Maria Rosaria Nucera, Laura Colombo, Saverio Paltrinieri, Renzo Bagnati, Jason Thomas Duskey, Riccardo Caraffi, Maria Angela Vandelli, Franco Taroni, Mario Salmona, Eugenio Scanziani, Gerardo Biella, Barbara Ruozi, Giovanni Tosi, Elena Cattaneo

## Abstract

Evidence that Huntington’s disease (HD) is characterized by impaired cholesterol biosynthesis in the brain has led to strategies to increase its level in the brain of the rapidly progressing R6/2 mouse model, with a positive therapeutic outcome. Here we tested the long-term efficacy of chronic administration of cholesterol to the brain of the slowly progressing zQ175DN knock-in HD mice in preventing (“early treatment”) or reversing (“late treatment”) HD symptoms. To do this we used the most advanced formulation of cholesterol loaded brain-permeable nanoparticles (NPs), termed hybrid-g7-NPs-chol, which were injected intraperitoneally.

We show that one cycle of treatment with hybrid-g7-NPs-chol, administered in the presymptomatic (“early treatment”) or symptomatic (“late treatment”) stages is sufficient to normalize cognitive defects up to 5 months, as well as to improve other behavioral and neuropathological parameters. A multiple cycle treatment combining both early and late treatments (“2 cycle treatment”) lasting 6 months generates therapeutic effects for more than 11 months, without severe adverse reactions.

Sustained cholesterol delivery to the brain of zQ175DN mice also reduces mutant Huntingtin aggregates in both the striatum and cortex and completely normalizes synaptic communication in the striatal medium spiny neurons compared to saline-treated HD mice. Furthermore, through a meta-analysis of published and current data, we demonstrated the power of hybrid-g7-NPs-chol and other strategies able to increase brain cholesterol biosynthesis, to reverse cognitive decline and counteract the formation of mutant Huntingtin aggregates.

These results demonstrate that cholesterol delivery via brain-permeable NPs is a therapeutic option to sustainably reverse HD-related behavioral decline and neuropathological signs over time, highlighting the therapeutic potential of cholesterol-based strategies in HD patients.

## Introduction

A growing number of studies highlight the importance of cholesterol homeostasis for brain function. Disruption of its synthesis and/or catabolism is associated with several neurological disorders (Martin *et al*, 2014). Among them is Huntington’s disease (HD), an autosomal dominant neurodegenerative disease whose symptoms typically occur in midlife and caused by the expansion of a polyglutamine-encoding cytosine adenine guanine (CAG) tract in exon 1 of the huntingtin (HTT) gene (Saudou & Humbert, 2016). Due to this mutation, the striatal medium spiny neurons (MSNs) and cortical neurons projecting to the striatum progressively degenerate, causing motor defects, cognitive decline, and psychiatric disturbance (Zuccato *et al*, 2010; Rub *et* al, 2016).

Among the underlying pathogenetic mechanisms, evidence accumulated over the past two decades has implicated impaired cholesterol biosynthesis in the brain of HD rodent models, with early and massive involvement of the striatum (Valenza *et al*, 2007a; Valenza *et al*, 2007b; Kacher *et al*, 2019). In particular, reduced levels of cholesterol precursors and *de novo* cholesterol synthesis were measured in the brain of several rodent models of HD from presymptomatic stages (Valenza *et al*, 2007a; Valenza *et al*, 2007b; Shankaran *et al*, 2017). Because peripheral cholesterol is unable to cross the blood-brain barrier (BBB) (Jurevics & Morrel, 1995) and local synthesis of cholesterol in the brain is critical for neuronal function and synaptic transmission (Bjorkhem *et al*, 2004; Li *et al*, 2022), its reduced biosynthesis may contribute to the severe cognitive and synaptic defects observed in the disease (Zuccato *et* al, 2010; Rub *et al*, 2016). Levels of 24-S-hydroxycholesterol (24S-OHC), the brain-specific cholesterol catabolite which, unlike cholesterol, crosses the BBB and can be detected in plasma, were also decreased in the plasma of HD rodents (Valenza *et al*, 2007a) as in patients (Leoni *et al*, 2008; Leoni *et al*, 2011; Leoni *et al*, 2013). This finding is consistent with a defect in cholesterol biosynthesis in the brain, although it may also reflect ongoing neuronal loss (Leoni *et al*, 2008).

On this basis, we have previously begun to explore means of restoring cholesterol bioavailability in the brain of R6/2 mice, an HD mouse line that exhibits early and rapid progression of behavioural, molecular, and electrophysiological abnormalities, starting at 6 weeks of age. In one study, cholesterol loaded brain-permeable polymeric nanoparticles (NPs) made of poly-lactic-co-glycolic acid (PLGA) and modified on their surface with the endogenous g7 shuttle-peptide for brain penetration (Tosi *et al*, 2007), and called g7-PLGA-NPs-chol, were used to allow exogenous cholesterol to pass through the BBB after intraperitoneal (ip) injection. This strategy provided approximately 15 µg of cholesterol, which was sufficient to prevent cognitive decline and ameliorate synaptic dysfunction in R6/2 mice. However, motor behavior was not significantly improved (Valenza *et al*, 2015a).

Subsequent dose-dependent studies using osmotic minipumps allowed to infuse three increasing doses of cholesterol directly into the striatum of R6/2 mice in a constant and continuous manner (Birolini *et al*, 2020). The highest dose of 369 µg of cholesterol infused in 4 weeks was identified as the dose capable of restoring both cognitive and motor abnormalities, while the doses of 15 and 185 µg prevented cognitive decline without any benefit on motor performance.

Mechanistically, exogenous cholesterol normalized the synaptic activity of MSNs by increasing the number of glutamatergic synapses and the number of docked GABAergic synaptic vesicles and counteracted the aggregation of mutant HTT (muHTT) by modulating lysosome-dependent pathways (Birolini *et al*, 2020). Restoration of synaptic transmission and clearance of muHTT aggregates were also confirmed after injection into the striatum of R6/2 mice of a recombinant AAV carrying the SREBP2 transcription factor known to activate the expression of most cholesterol biosynthesis genes (Birolini *et al*, 2021a). In addition to increasing the transcription of genes of the cholesterol biosynthesis pathway, exogenous SREBP2 also restored dopamine receptor D2 transcript levels in the R6/2 striatum (Birolini *et al*, 2020), indicating an overall improvement of several molecular, cellular, and biochemical parameters in animals exposed to brain cholesterol-raising strategies.

A more advanced formulation of NPs containing 30-times more cholesterol than g7-PLGA-NPs-chol was recently developed (Belletti *et al*, 2018). These NPs, named hybrid-g7-NPs-chol because g7-PLGA is combined with cholesterol itself in the formulation, completely rescued cognitive dysfunction. However, they partially ameliorated the motor abnormalities in R6/2 mice (Birolini *et al*, 2021b) due to the rapid progression and short life span (13 weeks) of this HD mouse model and the fact that not all of cholesterol is released from the injected NPs in this short period of time (Birolini *et al*, 2021b), preventing the full exploitation of the benefits of the treatment.

Here, we tested the long-term effects of hybrid-g7-NPs-chol on disease progression in the slow-progressing zQ175DN (delta-neo) heterozygous (het) knock-in mice (Menalled *et al*, 2012; Southwell *et al*, 2016). This mouse model more closely mimics the human condition as the neo-deleted knock-in allele encoding the human HTT exon 1 sequence with a CAG repeat stretch of ∼190 is inserted into the mouse huntingtin (htt) gene. Mice show cognitive and motor deficits starting from 6 and 9 months, respectively, while clinical signs and survival deficits appear at 12 months of age (Southwell *et al*, 2016).

In our study we analyzed the effect of the treatment over 1 year, from the prodromal stage to the symptomatic stage. The prolonged survival of the mice ensured complete release of cholesterol from the injected NPs. In a first cohort, animals were exposed to a single cycle of treatment (2 ip injections/week for 5 weeks) in the presymptomatic phase (“early treatment”) to evaluate its influence on the onset of the disease and its efficacy over time. In a second cohort, animals were exposed to a single cycle of treatment when symptoms were evident (“late treatment”) to test cholesterol effectiveness in restoring physiological conditions. The last cohort included animals doubly treated, i.e. with “early treatment” and “late treatment” (“2-cycle treatment”) to simulate long-term multiple dose treatment. Improvement of disease phenotypes were analyzed along with the likelihood of side effects. In all three studies, cholesterol administration led to full recovery from the cognitive decline associated with disease progression, an effect that was long-lasting and occurred even with the “late treatment” alone, when the animals were symptomatic. Reversal of motor defects and massive reduction of muHTT aggregates coincided with the timing of maximum cholesterol release. No major tissue or systemic side effects were noted.

We conclude that cognitive decline in HD mice is fully curable for at least 9 consecutive months and that cholesterol delivery to the HD brain is a safe and versatile therapeutic option that can prevent or counter multiple disease phenotypes.

## Results

### *In vivo* distribution and pharmacokinetics of hybrid-g7-NPs-chol

To design the therapeutic regimens for NP-based cholesterol delivery to zQ175DN mice, we first traced the temporal dynamics of intracellular cholesterol release from NPs using hybrid-g7-NPs labelled with Cy5 (a far-red fluorescent dye) and loaded with Bodipy-cholesterol, a photostable fluorescent cholesterol derivative. The coefficient of overlap between the Cy5 (detecting the NPs) and Bodipy (tracking the cholesterol moiety) signals measured by confocal microscopy was then used to assess cholesterol release from NPs over time.

7-week-old wt mice were treated with a single ip injection, sacrificed at 24 hours, 2 weeks, 10 weeks, and 20 weeks post-injection and overlap coefficient was measured over time on the cryo-sectioned brain slices (**Fig EV1A**). We found that 24 hours after a single ip injection, Cy5 and Bodipy signals colocalized in striatum and cortex (overlap coefficient = 0.829 and 0.758 respectively; an overlap coefficient of 1 indicates total overlap), while 2 weeks after the two signals were partially separated (**Fig EV1B and C**). Overlap coefficient at 2 weeks showed that approximately 15% and 28% of Bodipy were no longer colocalized with Cy5 in striatum and cortex, respectively, suggesting a progressive release of cholesterol from the NPs, in parallel with a reduction in Cy5 signal likely due to polymer degradation (overlap coefficient = 0.708 in striatum and 0.542 in cortex; **Fig EV1B and C**). At 10 weeks post injection, 70% of cholesterol was released in the striatum and 63% in the cortex (overlap coefficient = 0.232 and 0.274, respectively), while at 20 weeks 93% of cholesterol was released in the striatum and 86% in the cortex (overlap coefficient = 0.057 and 0.109 respectively; **Fig EV1B and C**). Collectively, these data show that, in the brain, all encapsulated cholesterol is released within 20 weeks.

To further strengthen our understanding of NPs pharmacokinetics and quantify the amount of exogenous cholesterol reaching the brain, we then exploited deuterated cholesterol (d6-chol)-laden hybrid-g7-NPs (hybrid-g7-NPs-d6-chol) (**Fig EV1D**). The presence of six deuterium atoms in the cholesterol structure makes it possible to distinguish exogenous from endogenous cholesterol and to quantify the former in different tissues by mass spectrometry (Birolini *et al*, 2021b). Hybrid-g7-NPs-chol carried a calculated amount of about 29 mg of cholesterol for 100 mg of NPs (see Methods). In our previous study we demonstrated that the concentration of d6-chol in different brain regions (striatum, cortex, cerebellum) at 24 hours ranged from 0.36 to 0.54 ng/mg of tissue, with values increasing up to three times at the 2-week time point (Birolini *et al*, 2021b) (summarized in **Table EV1**). We have now added the time points 10- and 20-week post ip injection (red boxes in **Table EV1**). D6-chol was still present in the brain at 10 weeks, although the amount decreased by half from 2 weeks, and was further reduced at 20 weeks before returning to levels seen 24 hours after treatment. Importantly, d6-chol in peripheral tissues and plasma decreased rapidly over time and was no longer detected at the late time points, suggesting that cholesterol was utilized by brain cells and cleared in the periphery (**Table EV1**).

### One cycle of treatment with hybrid-g7-NPs-chol in the presymptomatic phase postpones symptoms in zQ175DN mice

In the first experimental regimen paradigm, defined “early treatment”, zQ175DN mice were ip injected with hybrid-g7-NPs-chol (here referred to as the “zQ175DN+chol” group) at presymptomatic stage from 5 to 9 weeks of age, following the same regimen used for R6/2 mice, i.e. with 2 injection/week for 5 weeks (Birolini *et al*, 2021b). As controls, wt and zQ175DN mice were treated with saline (herein named wt and zQ175DN, respectively). Since about 493 µg of cholesterol were administered with each ip injection (see methods), a cycle of 10 ip injections corresponded to a total of about 4930 µg of cholesterol administered to each animal. This amount was about 30 times higher than the amount used in our previous study with g7-PLGA-NPs-chol injected in R6/2 mice (Valenza *et al*, 2015a). Since 10% of g7-NPs cross the BBB (Tosi *et al*, 2007; Tosi *et al*, 2014), 493 µg of cholesterol is expected to reach the brain per treatment cycle, a dose similar to the higher dose delivered into the striatum with osmotic mini-pumps and found to improve both cognitive and motor defects in R6/2 mice (Birolini *et al*, 2020).

Cognitive and motor skills were extensively assessed in each group at 20, 29, 35 and 45-48 weeks of age (**Fig 1A; Table EV2**).

**Figure 1.**
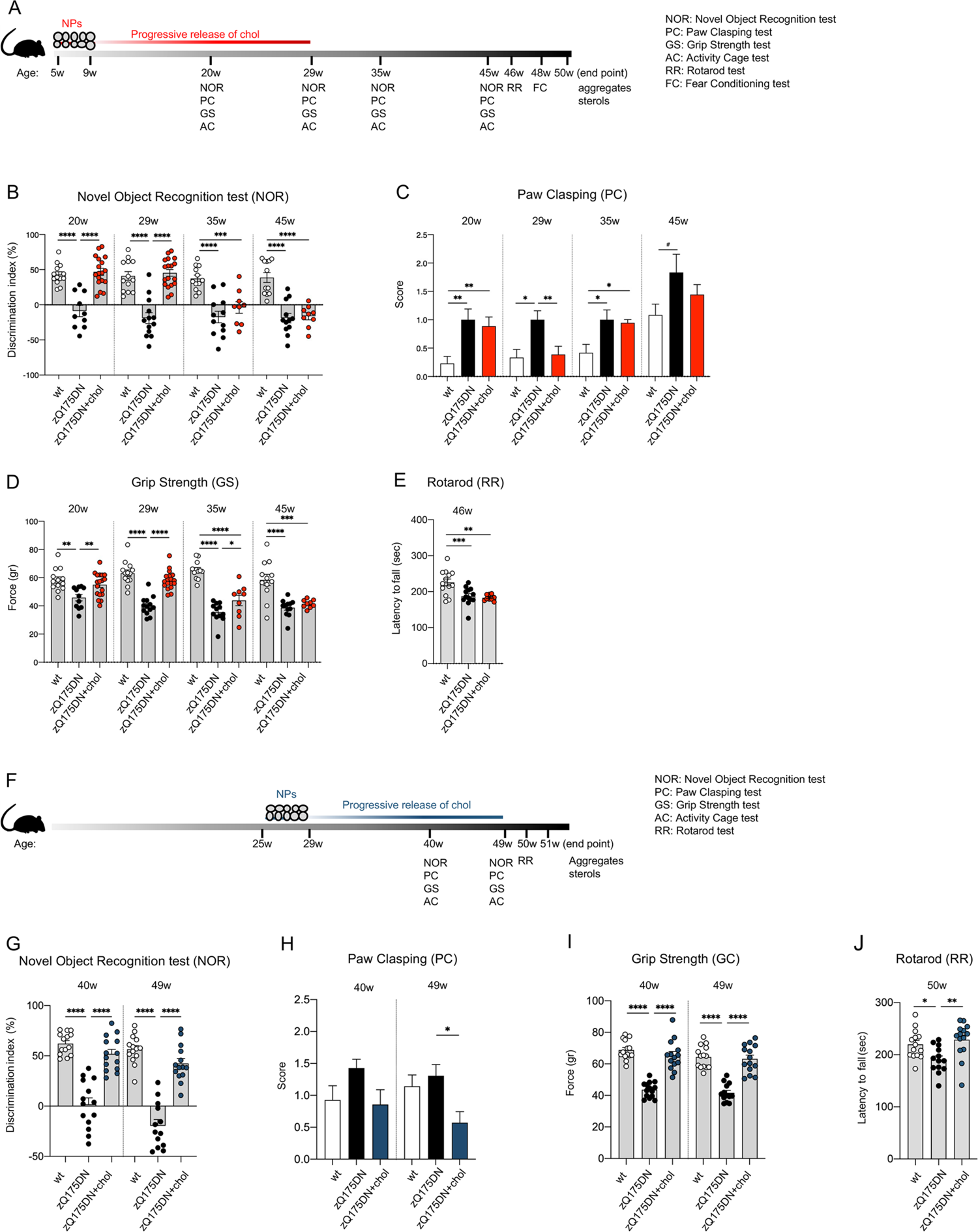
Cognitive and motor abilities of zQ175DN mice after “early treatment” and “late treatment”. **A** Experimental paradigm of the “early treatment”: zQ175DN mice (*N* = 9-18) were treated with g7-hybrid-NPs-chol (0.12 mg NPs/gr body weight) from 5 to 9 weeks of age with 2 ip injection/week; wt (*N* = 10-13) and zQ175DN (*N* = 10-11) littermates were treated with saline solution as controls. Novel Object Recognition (NOR), Paw Clasping (PC), Grip Strength (GS), and Activity Cage (AC) tests were performed at 20-29-35-45 weeks of age, Rotarod (RR) at 46 weeks of age, Fear Conditioning (FC) at 48 weeks of age. Mice were sacrificed at 50 weeks of age. **B-E** Behavioral tests. Discrimination index (DI; %) in NOR (B); PC score (C); GS (gr, D); latency to fall (sec) in RR (E). **F** Experimental paradigm of the “late treatment”: zQ175DN mice (N = 13-14) were treated with g7-hybrid-NPs-chol (0.12 mg NPs/gr body weight) from 21 weeks of age to 25 weeks of age with 2 ip injection/week; wt (N = 13-14) and zQ175DN (N = 13-14) littermates were treated with saline solution as controls. NOR, PC, GS, and AC were performed at 40-49 weeks of age, RR at 50 weeks of age, and mice were sacrificed at 51 weeks of age. **G-J** Behavioral tests. Discrimination index (DI; %) in NOR (G); PC score (H); GS (gr, I); latency to fall (sec) in RR (J). (C and H): wt mice develop clasping with age; the mean of combined PC data in wt mice at 40, 45 and 49w (C and H) is higher than the mean of PC data in wt mice at 20, 29, 35w (C) (1.034 +/− 0,117 and 0,2892 +/− 0.05 respectively; p< 0.0001 Welch’ t test). Data information: data in B-E are from three independent trials and shown as scatterplot graphs with mean±SEM. Data in G–J are from two independent trials and shown as scatterplot graphs with mean±SEM. Each dot (B, D, E, G, I, J) corresponds to the value obtained from each animal. Statistics: one-way ANOVA with Tuckey post-hoc test (*p<0.05; **p<0.01; ***p<0.001; ****p<0.0001).

To evaluate cognitive performance after treatment, we first performed the Novel Object Recognition (NOR) test, which measures recognition memory, a subtype of declarative long-term memory, and is based on rodents’ natural propensity to explore novelty. Recognition memory was severely impaired in zQ175DN mice as early as 20 weeks of age and the defect persisted up to the 45-week time point tested (negative Discrimination Index in **Fig 1B**). After the “early treatment”, zQ175DN mice behaved like the wt mice at 20 and 29 weeks of age but the effect was lost at 35 and 45 weeks (**Fig 1B**). This finding is consistent with the kinetics of cholesterol release, accumulation, use and elimination in the brain as described in **Figure EV1** and in **Table EV1** and demonstrates the ability of exogenous cholesterol to postpone the onset of abnormal cognitive events by several weeks in zQ175DN mice.

To evaluate the impact of exogenous cholesterol on disease progression, we also measured dystonic movements through the Paw Clasping (PC) test. zQ175DN mice exhibited clasping activity as early as 20 weeks of age, which was not rescued in zQ175DN+chol mice (**Fig 1C**). However, at 29 weeks of age, the amount of cholesterol released from the NPs was sufficient to normalize clasping behaviour. As for the NOR, this effect was lost at 35 and 45 weeks of age (**Fig 1C**).

In the Grip Strength (GS) test, which measures neuromuscular strength, zQ175DN mice showed a defect in this parameter compared to controls at all time points (**Fig 1D**). Administration of cholesterol to the brain completely reversed this defect at 20 and 29 weeks of age. At 35 weeks the improvement was markedly diminished although still significant and was lost at 45 weeks (**Fig 1D**).

At 46 weeks of age, Rotarod (RR) test was used to assess motor skills. A decrease in the latency to fall was observed in zQ175DN mice compared to wt littermates, confirming the late-onset motor defect observed in this mouse model (Menalled *et al*, 2012). However, not unexpectedly, this late defect was not recovered by the “early treatment” (**Fig 1E**).

Similarly, at 48 weeks of age, there was no significant effect of the early cholesterol treatment in the Fear Conditioning (FC) test (**Fig EV2A and B**), a type of associative learning task that measures the animal’s ability to associating a neutral conditional stimulus (tone) with an unconditioned aversive stimulus (a mild electrical foot shock) exhibiting a conditioned fear response resulting in no movement (freezing). zQ175DN mice showed reduced freezing in both the contextual (when the animal is tested in the original training context) and cued (when the conditional stimulus is presented in a different context) paradigm, but as for the GS (**Fig 1D**) and RR (**Fig 1E**) tests performed at 45-46 weeks, the “early treatment” had no impact on FC measured at the late 48-week time point (**Fig EV2A and B**).

These data indicate that the “early treatment” prevents disease manifestations and postpones the onset of symptoms to week 29 (as per NOR and PC tests) or week 35 (as per GS tests) in treated animals, i.e. 20-26 weeks (5-6.5 months) after the last ip injection. Not unexpectedly, the “early treatment” produced no measurable effects at 48 weeks (11 months) in all tests.

### Treatment is also effective in the symptomatic phase

In a second experimental regimen paradigm, we tested the ability of hybrid-g7-NPs-chol to counteract disease manifestations in symptomatic zQ175DN mice treated from 25 to 29 weeks of age (termed “late treatment”) (**Fig 1F**). Unlike the “early treatment” (**Fig 1B**), zQ175DN+chol mice showed a complete recovery of NOR-dependent cognitive decline at 40 and 49 weeks of age, compared with saline-treated zQ175DN mice (**Fig 1G**).

Regarding the PC test, we found that at 49 weeks of age, zQ175DN+chol mice performed better than wt and zQ175DN groups (**Fig 1H**). The latter two groups did not differ from each other as the wt mice also developed clasping with age.

In contrast, neuromuscular defects were fully rescued at both 40 and 49 weeks of age compared to untreated zQ175DN mice (**Fig 1I**), i.e. at the late time points that showed no rescue in the “early treatment” (**Fig 1D**). Finally, while cholesterol administration at presymptomatic stage was not effective in the RR test (**Fig 1E**), zQ175DN mice from the “late treatment” group showed improved performance at 50 weeks of age (**Fig 1J**).

### The combination of “early and late” cholesterol administration leads to complete cognitive and motor recovery

The finding that the beneficial effect of one cycle of treatment persisted at 29 weeks of age (i.e. 20 weeks after the last ip injection, **Fig 1B-D**) but was lost starting at 35 weeks of age, suggested that more treatments can prolong the benefit over time. Consequently, we tested the combination of two treatment cycles, which we termed “2-cycle treatment” (**Fig 2A**): the first from 5 to 9 weeks of age (as performed with the “early treatment”, **Fig 1A**) and the second from 21 to 25 weeks of age, for a total of 20 injections per animal over 20 weeks (5 months).

**Figure 2.**
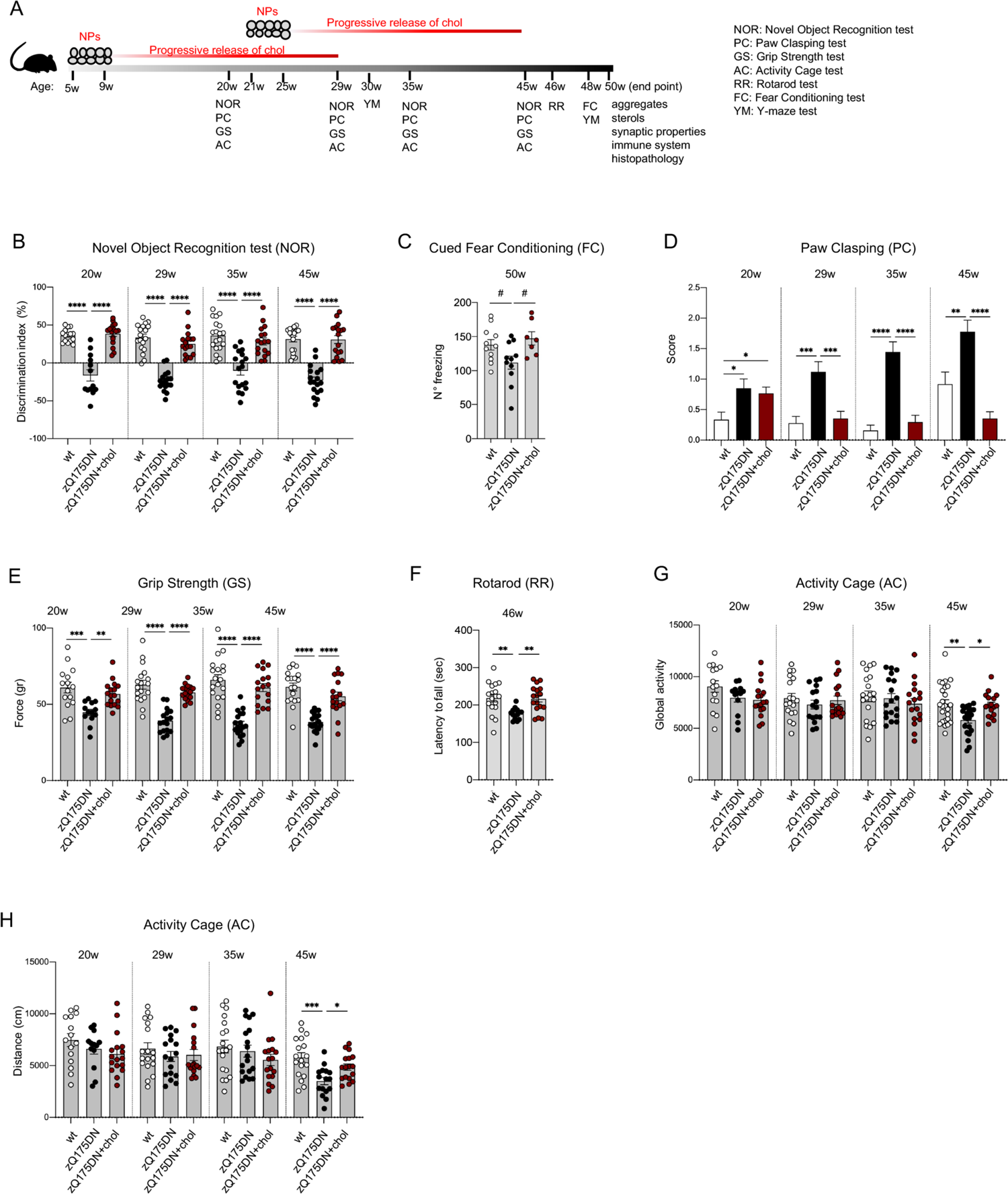
Cognitive and motor abilities of zQ175DN mice after “2-cycle treatment”. **A** Experimental paradigm of the “2-cycle treatment”: zQ175DN mice (*N* = 9-17) were treated with g7-hybrid-NPs-chol (0.12 mg NPs/gr body weight) from 5 weeks of age to 9 weeks of age and from 21 to 25 weeks of age with 2 ip injection/week/cycle; wt (*N* = 15-21) and zQ175DN (*N* = 11-21) littermates were treated with saline solution as controls. NOR, PC, GS, and AC were performed at 20-29-35-45 weeks of age, Y-maze (YM) test at 30-48 weeks of age, RR at 46 weeks of age, FC at 48 weeks of age. Mice were sacrificed at 51 weeks of age. **B-H** Behavioral tests. Discrimination index (DI; %) in NOR (B); n° of freezing episodes in FC (cued paradigm, C); PC score (D); GS (gr, E); latency to fall (sec) in RR (F); global activity (G) and distance (cm, H) in AC. Data information: data in B–H are from three independent trials and shown as scatterplot graphs with mean±SEM. Each dot (B, D–H) corresponds to the value obtained from each animal. Statistics: one-way ANOVA with Tuckey post-hoc test (*p<0.05; **p<0.01; ***p<0.001; ****p<0.0001).

After “2-cycle treatment”, complete reversal of cognitive decline associated with recognition memory (NOR test) was observed at all analysed time points in zQ175DN mice, and up to the study endpoint of 45-50 weeks of age (**Fig 2B**). In the FC test, the same treatment regimen also restored normal freezing at 50 weeks in zQ175DN mice exposed to the cued paradigm (**Fig 2C**), although this was not true in the contextual paradigm (**Fig EV2C**). To further explore aspects of cognition, we also performed the Y-Maze (YM) spontaneous alternation test, which measures the rodents’ willingness to explore new environments and provides an index of their short-term memory. However, this test revealed no difference at 30 and 48 weeks between the genotypes (**Fig EV2D**).

Furthermore, disease progression assessed by the PC test was halted in zQ175DN+chol mice treated with 2 cycles starting at 29 weeks of age, and the improvement remained highly significant at 35 and 45 weeks of age (**Fig 2D**). At 45 weeks of age, zQ175DN+chol mice performed even better than wt mice, which worsen over time due to ageing (**Fig 1C and H, Fig 2D**). Thus, the “2-cycle treatment” ensures continued benefit in NOR and PC tests that extends to times when the “early treatment” alone was ineffective, due to depletion of the initially administered cholesterol over the course of several months (**Fig 1B and C**).

Also in the GS test, where the efficacy of “early treatment” was moderately present at 35 weeks of age and absent at 45 weeks of age (**Fig 1D**), “2-cycle treatment” produced a complete recovery even at the late time points, with values overlapping those of wt mice (**Fig 2E**).

Regarding motor defects, 2-cycle treatment with hybrid-g7-NPs-chol normalized RR performance at 46 weeks in zQ175DN+chol mice compared to saline-treated mice, reaching values similar to those observed in wt mice (**Fig 2F**). Regarding other motor skills (assessed by the Activity Cage test, AC), a small, but significant, reduction in global activity and total distance travelled was measured at 45 weeks of age in zQ175DN mice compared to wt mice and the 2-cycle treatment was able to counteract these defects (**Fig 2G and H**). These benefits were not seen in the zQ175DN+chol mice after the “early” or “late treatment” (**Fig EV2E-H**, see time points 45-49 weeks).

**Figure 3A** shows the heat-maps generated by aggregating the values of cognitive (NOR) and motor (PC, GS, AC) tasks from “early treatment”, “late treatment” and “2-cycle treatment” and using a color-coding system to represent different levels of performance (from best, light blue, to worst, dark blue, performance). As expected, zQ175DN mice globally deteriorated over time, while zQ175DN+chol mice after the “early treatment” performed better up to 29 weeks of age (i.e., for 20 weeks after the last ip injection) compared to untreated HD littermates (**Fig 3A**, top). Similarly, HD-related defects were recovered in the symptomatic HD mice after the “late treatment” (**Fig 3A**, center). Of note, a “2-cycle treatment” (**Fig 3A**, bottom) was required to have complete and sustained recovery at all time points.

**Figure 3.**
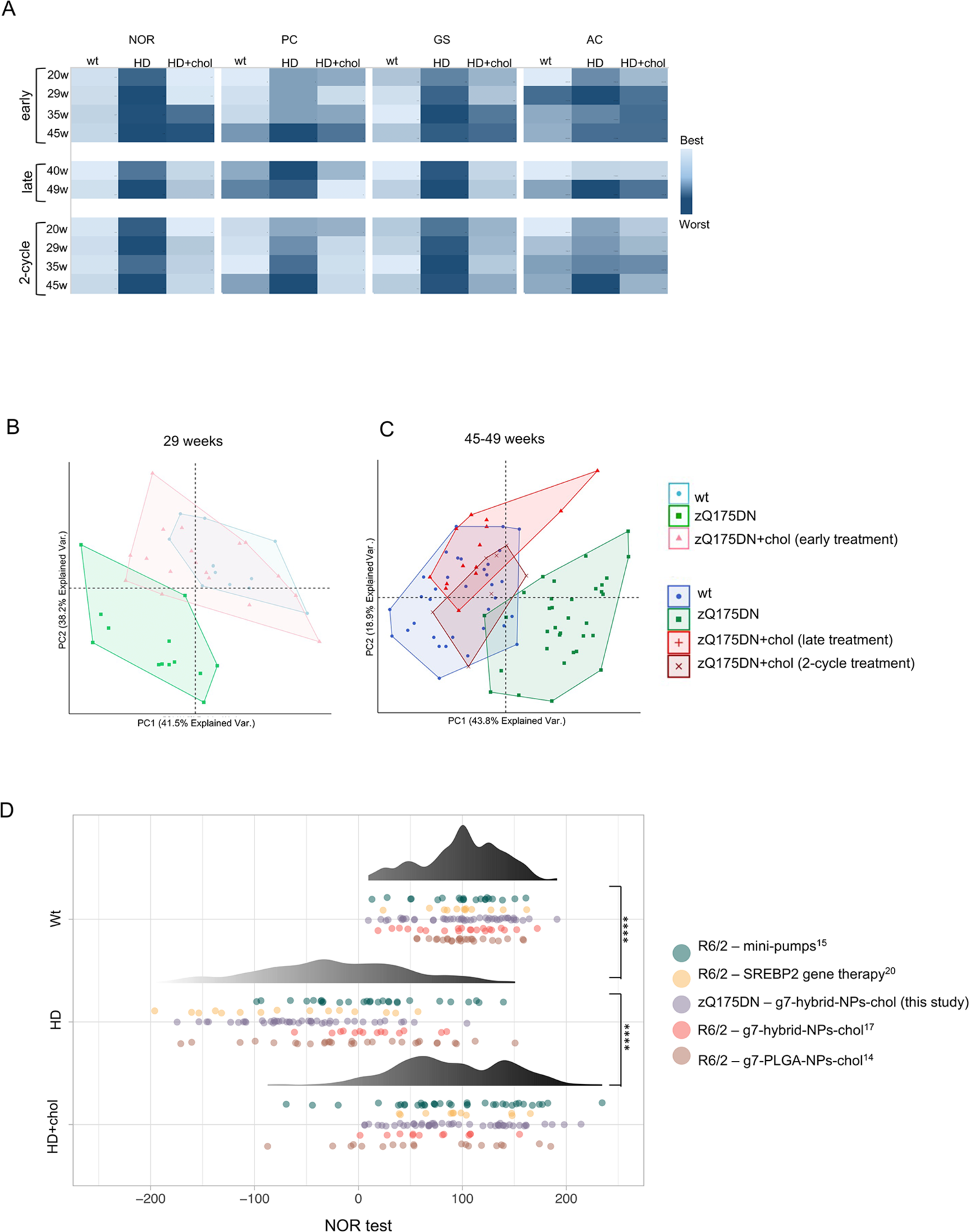
Segregation of the behavioral tasks in cholesterol treated HD mice with wt mice versus HD mice. **A** Heat maps summarizing the behavior related to NOR, PC, GS, and AC tests of all treatment using conditional formatting (excel). Light blue: best performance; dark blue: worst performance. **B-C** Principal component analysis (PCA) by combining all the values related to motor and cognitive tasks from all mice at 29 weeks (B) and 45-49 weeks of age (C). The convex hull of the set of points belonging to each group of mice was also visualized. **D** Data-visualization of the overall cognitive performance assessed by NOR test among wt, HD and HD cholesterol-treated mice, comprehensive of data collected from five different studies. Data were visualized using a dotplot also showing their distribution. The differences between groups have been assessed using a pairwise t-test, multiple testing correction was performed using the Bonferroni method.

Principal component analysis (PCA) applied to the values of all motor and cognitive tests also clearly separated the wt group from the zQ175DN group (**Fig 3B and C**). Importantly, at 29 weeks of age, after the “early treatment”, the zQ175DN+chol group separated from the zQ175DN group and showed overlap with the wt group (**Fig 3B**). Similarly, at 45-49 weeks of age, zQ175DN+chol mice after the “late treatment” or the “2-cycle treatment” were clearly distinguishable from the zQ175DN group and overlapped with the wt group (**Fig 3C**).

To explore further the efficacy of cholesterol administration on cognitive decline, we decided to pool and re-analyse all raw data obtained in the NOR assays performed on R6/2 mice (Birolini *et al*, 2021b) and zQ175DN mice (this paper) treated with hybrid-g7-NPs-chol, R6/2 mice treated with the previous g7-PLGA-NPs-chol formulation (Valenza *et al*, 2015a), as well as R6/2 mice implanted with osmotic mini-pumps filled with cholesterol (Birolini *et al*, 2020) or intracerebrally injected with AVV-SREBP2 (Birolini *et al*, 2021a). From all these studies only the latest time point was considered for a total of 124 wt (control) mice, 130 untreated HD mice and 127 treated HD mice (the last two groups included both R6/2 and zQ175DN mice) which were then statistically compared. As shown in **Fig 3D**, the distribution of the NOR values obtained from two HD mouse lines treated with any of the four treatments mentioned above, including those used in this paper, regardless of the type of strategy, delivery system, mouse genotype and time of administration, overlapped more with the values of the wt groups than with those of the HD groups. This result highlights that any treatment that supplies exogenous cholesterol to the brain or increases its endogenous synthesis is able to counteract cognitive decline in HD mice. In the present study we demonstrate that the administration of cholesterol to the brain leads to a cognitive recovery that lasts almost 1 year.

In this study, we also quantified in the striatum and cortex of saline- and chol-treated animals the content of total cholesterol and the level of its main precursors lanosterol, lathosterol, and desmosterol, and of the brain-specific cholesterol catabolite 24S-OHC at 29 and 50-51 weeks of age, i.e. 5-6 months after the last ip injection in the different experimental regimens adopted (**Fig 4**; **Fig EV3; Fig EV4**). In total, by combining all three trials, we performed 480 measurements of the level of cholesterol and its metabolites/catabolites in the striatum and cortex of 16 mice for each of the three groups (wt, zQ175DN and 16 zQ175DN+chol mice). We confirmed that a statistically significant reduction in the level of cholesterol, lanosterol, lathosterol and desmosterol is present in zQ175DN striatum at all time-points (29 and 50 weeks), compared to wt mice (**Fig EV3**). These data reinforce previous mass-spectrometry data showing reduced cholesterol content and cholesterol biosynthesis in the striatum of R6/2 mice and other HD mouse models (Valenza *et al*, 2007b; Birolini *et al*, 2021a) and a reduced rate of daily cholesterol synthesis *in vivo* in zQ175 mice carrying the floxed neo cassette (Shankaran *et al*, 2017). A decrease in 24S-OHC level was also found in the striatum of zQ175DN compared to wt mice, albeit with some variability between the trials (**Fig EV3**). A similar reduction in lathosterol, lanosterol and 24S-OHC was observed in cortical samples from zQ175DN mice (**Fig EV4).** Not unexpectedly, at the last time points analysed, i.e. 20 and 40 weeks after the last ip injection of hybrid-g7-NPs-chol, we found no differences in the steady-state levels of all the metabolites in treated versus untreated zQ175DN mice (**Fig 4**; **Fig EV3**; **Fig EV4**).

**Figure 4.**
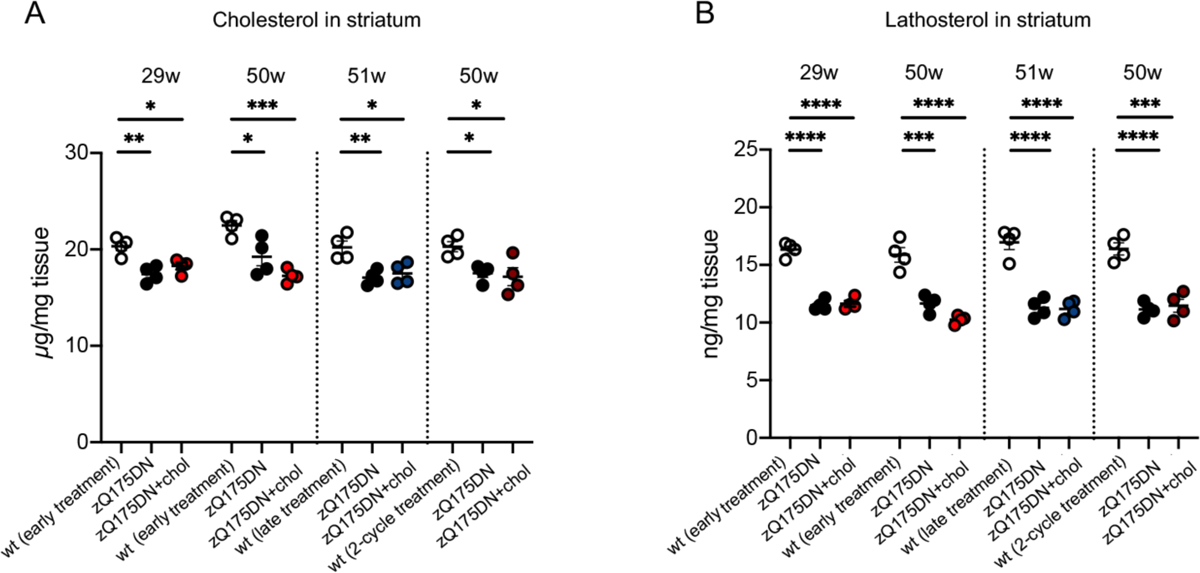
Sterols in the striatum of zQ175DN mice after “early treatment”, “late treatment” and “2-cycle treatment”. **A-B** Cholesterol (A) and lathosterol (B) content quantified by mass spectrometry in the striatum of animals (*N* = 4/group) from the “early treatment” (29- and 50-week), “late treatment” (51-week) and “2-cycle treatment” (50-week). Data information: data in A-B are shown as scatterplot graphs with mean±SEM. Each dot corresponds to the value obtained from each animal. Statistics: one-way ANOVA with Tuckey post-hoc test (*p<0.05; **p<0.01; ***p<0.001; ****p<0.0001).

Taken together, these data demonstrate that one cycle of hybrid-g7-NPs-chol, both in the presymptomatic and symptomatic phase, is sufficient to prevent or normalize several behavioral defects in zQ175DN mice for 5-6 months, while 2 cycles of hybrid-g7-NPs-chol generate global and long-lasting therapeutic effects that last almost a year.

### Exogenous cholesterol ameliorates the neuropathological deficit of HD mice

The evidence that exogenous cholesterol produces long-lasting benefits in zQ175DN mice led us to investigate whether early and late neuropathological abnormalities typical of the disease state were also ameliorated. Our previous work demonstrated that 4-week infusion of 369 µg of cholesterol with osmotic minipumps into the striatum of R6/2 mice reduced the number and size of muHTT aggregates (Birolini *et al*, 2020).

At 29 and 50 weeks of age, zQ175DN mice showed muHTT aggregates detected using EM48 antibody (**Fig 5**). In contrast, zQ175DN+chol mice that received one cycle of hybrid-g7-NPs-chol in the presymptomatic phase (“early treatment”; **Fig 5A**) showed a statistically significant reduction in the number of muHTT aggregates in both striatum and cortex at 29 weeks of age, i.e. 5 months after the last ip injection, but – consistently with the behavioural data in Figure 1 – this effect was lost at 50 weeks (**Fig 5B and C; Fig 5E and F**). In the cortex (**Fig 5G**), where more NPs are known to accumulate (Birolini *et al*, 2021b), but not in striatum (**Fig 5D**), the size of muHTT aggregates was also reduced at 29 weeks in cholesterol treated animals, an effect that also in this case was lost in the 50-week samples. Instead, at the same late time point, zQ175DN mice after the “late treatment” (**Fig 5H**) showed a significant reduction in the number and size of muHTT aggregates in both the striatum and cortex (**Fig 5I-N**). These data indicate that exogenous cholesterol promotes the clearance of muHTT aggregates rather than preventing their formation. Of note, the 50-week-old zQ175DN+chol mice that received 2 cycles of hybrid-g7-NPs-chol (**Fig 5O**) showed greater reductions in the number and size of muHTT aggregates in both striatum and cortex compared to zQ175DN control mice (**Fig 5P-U**).

**Figure 5.**
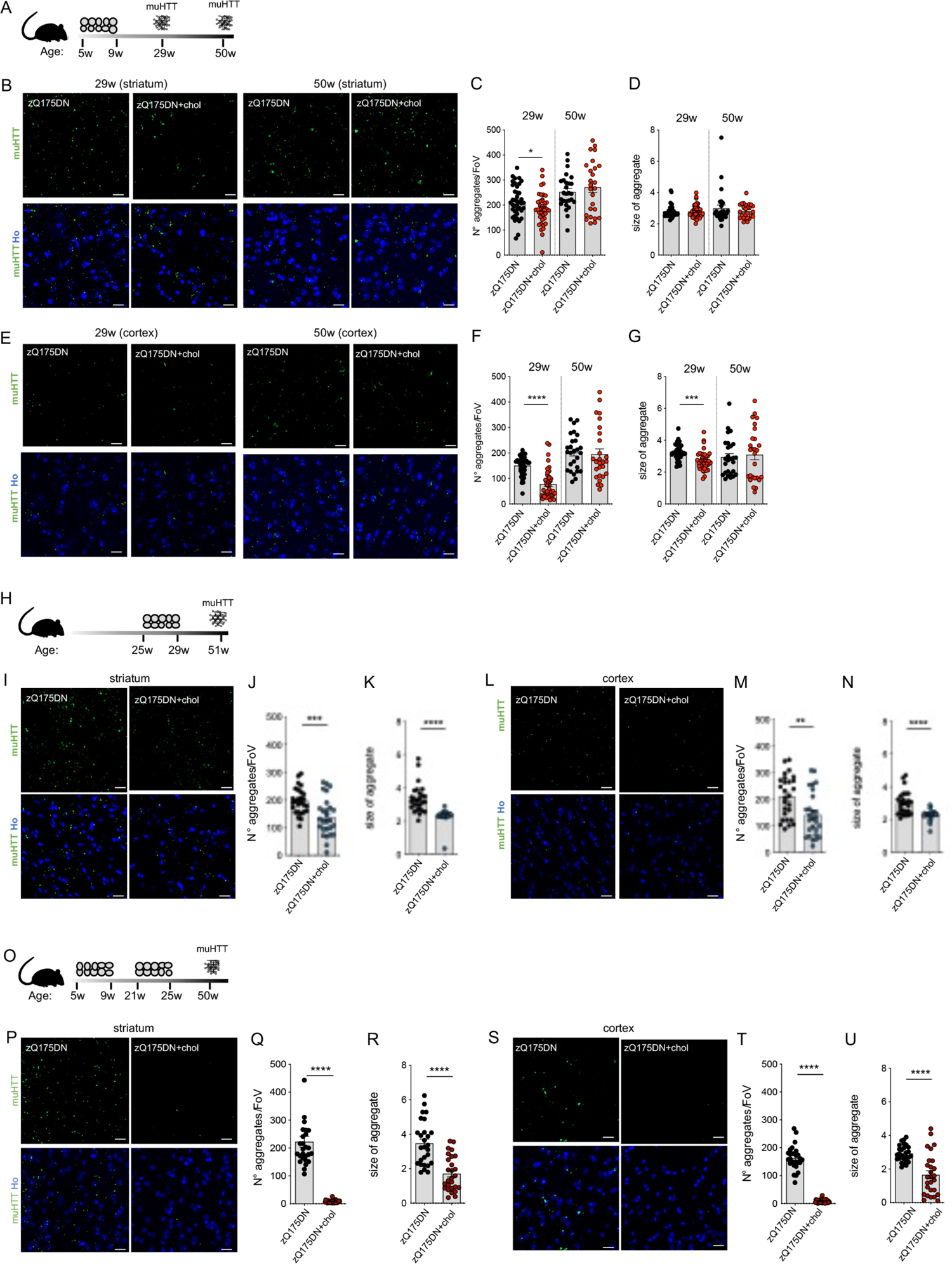
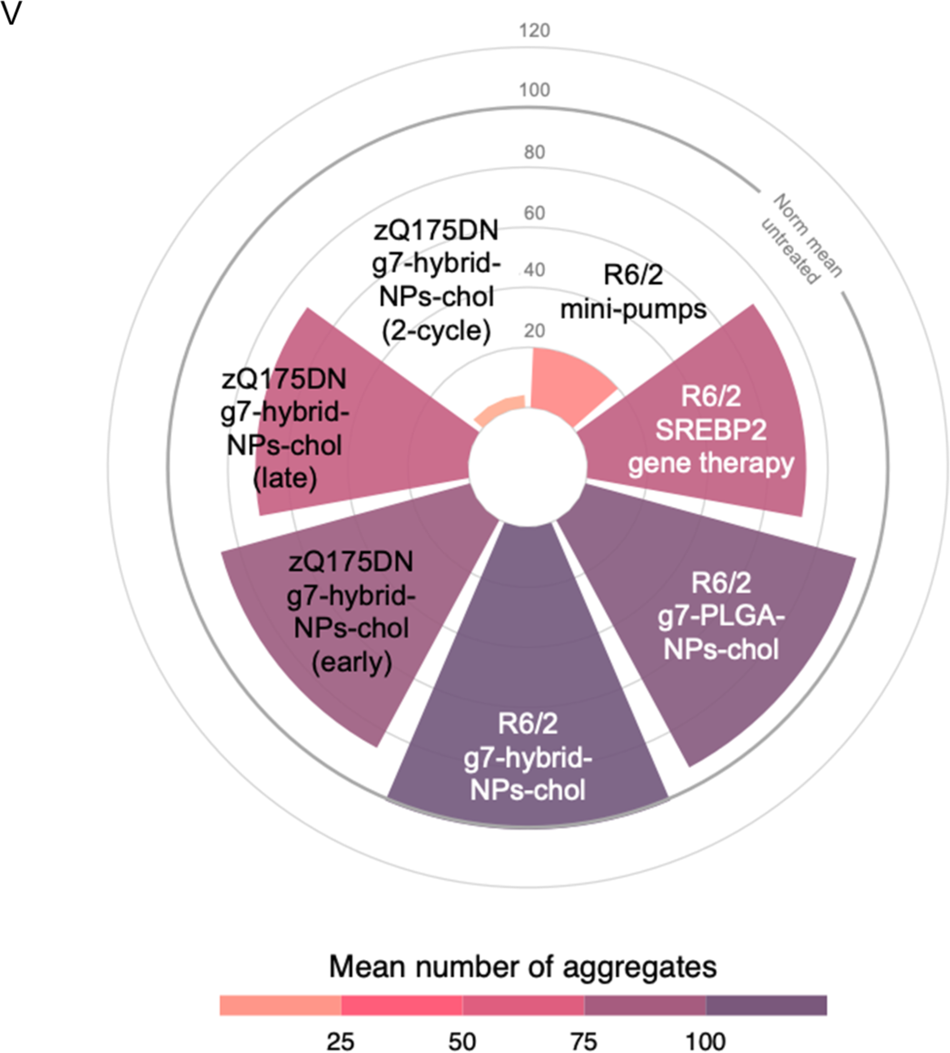
Mutant HTT aggregates in the striatum and cortex of zQ175DN mice after “early treatment”, “late treatment” and “2-cycle treatment”. **A–G** Experimental scheme of the “early treatment” (A) and immunolabeling of muHTT aggregates (showing muHTT aggregates positive for EM48 antibody in green) in striatum (B) and cortex (E) in brain coronal slices of zQ175DN and zQ175DN+chol mice at 29 and 50 weeks of age (*N* = 3/group); relative quantification of aggregates n° and size (C, D, F, G). **H-N** Experimental scheme of the “late treatment” (H) and immunolabeling of muHTT aggregates (showing muHTT aggregates positive for EM48 antibody in green) in striatum (I) and cortex (L) in brain coronal slices of zQ175DN and zQ175DN+chol mice at 51 weeks of age (*N* = 3/group); relative quantification of aggregates n° and size (J, K, M, N). **O-U** Experimental scheme of the “2-cycle treatment” (O) and immunolabeling of muHTT aggregates (showing muHTT aggregates positive for EM48 antibody in green) in striatum (P) and cortex (S) in brain coronal slices of zQ175DN and zQ175DN+chol mice at 50 weeks of age (*N* = 3/group); relative quantification of aggregates n° and size (Q, R, T, U). **V** Meta-analysis and data-visualization representing the mean of the number of muHTT aggregates in the striatum of HD mice after cholesterol-raising strategies. The normalized data were represented using a circular barplot to show the mean number of aggregates in the different treatments. Data information: Hoechst (Ho, blue) was used to counterstain nuclei. Scale bar: 20 µm. 10 images/animal were analyzed from 9 sections throughout the entire striatum and cortex. Data in C-D, F-G, J-K, M-N, Q-R, T-U are shown as scatterplot graphs with mean±SEM. Each dot corresponds to the value obtained from each image. Statistics: Student’s t-test (**p<0.01;***p<0.001; ****p<0.0001).

All these results demonstrate that cholesterol delivery to the HD brain through systemic injection of hybrid-g7-NPs-chol effectively reduces muHTT aggregation, and that the content and timing of the treatment are critical to detect changes.

We also collected and re-analysed all raw data on the number of muHTT aggregates in the striatum upon the different cholesterol-based strategies from our previous (Birolini *et al*, 2020; Birolini *et al*, 2021a; Birolini *et al*, 2021b) and current study. A total of 22 untreated and 27 treated zQ175DN mice were considered in this meta-analysis, for a total of 664 images collected for each group covering the striatum. The mean values for each group were normalized to 100% (**Table EV3**) and used to generate a circular barplot to visualize the effect on the number of muHTT aggregates upon the different treatments and strategies at the time points where a behavioural effect was observed. As shown in **Fig 5V**, all cholesterol-raising strategies were able to reduce muHTT aggregates but with varied degree. Treatment with either g7-PLGA-NPs-chol or g7-hybrid-NPs-chol in the R6/2 mouse model was ineffective as its short lifespan likely does not allow to benefit from the complete release of cholesterol from NPs. SREBP2 gene therapy (Birolini *et al*, 2021a), intrastriatal infusion of cholesterol with osmotic mini-pumps (Birolini *et al*, 2020) and the “2-cycle treatment” described in this paper were the most effective, with the latter having produced an almost complete disappearance of muHTT aggregates compared with any other strategy (**Fig 5V**).

### Neuronal functionality is restored by "2-cycle treatment”

Excitatory synaptic transmission is known to be severely affected in MSNs from several HD mouse models (Cepeda *et al*, 2006; Milnerwood & Raymond, 2010; Heikkinen *et al*, 2012). Increased input resistance (Rin) and decreased frequency of spontaneous excitatory post-synaptic currents (sEPSC) were described in MSNs of Q175 mice carrying the neo cassette compared to MSNs of wt mice (Milnerwood & Raymond, 2010; Heikkinen *et al*, 2012; Indersmitten *et al*, 2015; Vezzoli *et al*, 2019).

Here we examined the electrophysiological properties of striatal MSNs from brain slices of wt and zQ175DN mice versus those of zQ175DN+chol mice following the “2-cycle treatment”. The analysis of the basic membrane properties of MSNs did not show differences in membrane capacitance (Cm) between genotypes (**Fig 6A**). Instead, the resting membrane potential (Vm) was significantly depolarized, and membrane Input Resistance (Rin) was increased in zQ175DN MSNs with respect to wt MSNs (**Fig 6B and C**). Notably, cholesterol delivery normalized Vm in zQ175DN+chol MSNs bringing the value back to those of wt MSNs (**Fig 6B**).

**Figure 6.**
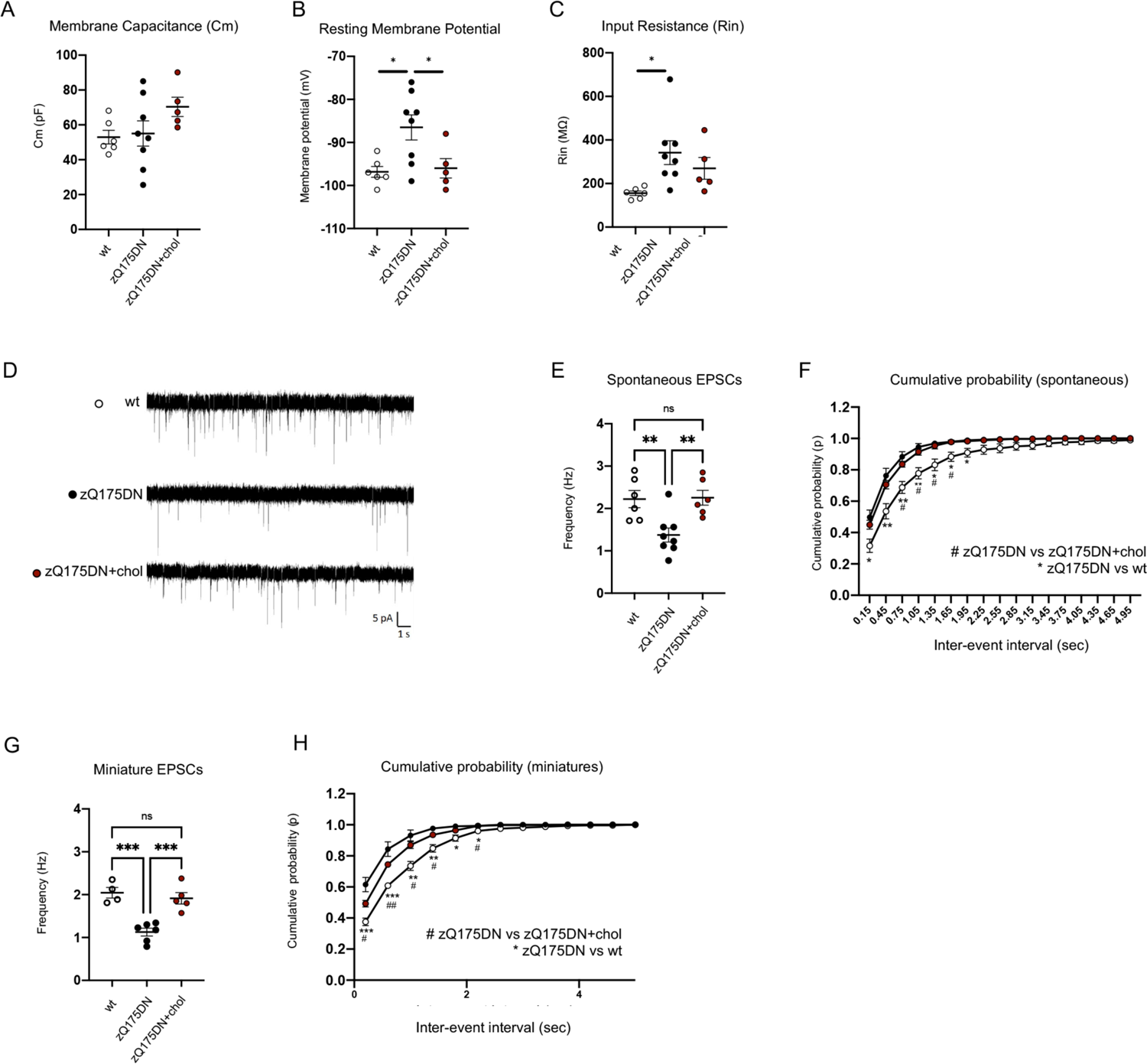
Electrophysiological analysis in MSNs of zQ175DN mice after “2-cycle treatment”. **A–C** Membrane Capacitance (Cm, A), Resting Potential (mV; B) and Input Resistance (Rin, C) recorded from MSNs of wt (6 cells from *N* = 3 mice), zQ175DN (8 cells from *N* = 4 mice) and zQ175DN+chol mice (5 cells from *N* = 3 mice) from the “2-cycle treatment”. **D** Representative traces of spontaneous EPSCs (sEPSCs) at a holding potential of −70 mV. **E–F** Mean frequency (E) and cumulative inter-event histogram (F) of spontaneous EPSCs (sEPSCs) recorded from MSNs of wt, zQ175DN and zQ175DN+chol mice. **G–H** Mean frequency (G) and cumulative inter-event histogram (H) of miniature EPSCs (mEPSCs) recorded from MSNs of wt, zQ175DN and zQ175DN+chol mice. Data information: data in A, B, C, E, and F are shown as scatterplot graphs with means±standard error. Each dot corresponds to the value obtained from each cell. Statistics (A-H): one-way ANOVA with Tuckey post-hoc test (*p<0.05; **p<0.01; ***P<0,001).

Furthermore, **Fig 6D and Fig 6E** show the representative traces and reduced frequency of spontaneous AMPA-mediated excitatory postsynaptic currents (sEPSC) in zQ175DN MSNs compared to wt and zQ175DN+chol MSNs. The frequency of sEPSC was fully normalized in cholesterol treated zQ175DN MSNs compared to zQ175DN MSNs (**Fig 6E).** Moreover, the distribution of the inter-event intervals of sEPSCs in the zQ175DN+chol MSNs overlapped with that of wt MSNs (**Fig 6F**). Similarly, the frequency and the inter-event intervals of miniature EPSCs (mEPSC), which are both altered in zQ175DN MSNs, were fully normalized in MSNs from zQ175DN+chol mice (**Fig 6G-H).** No significant differences were found in the mean amplitude and kinetics (rise time and decay time) of sEPSC (**Fig EV5A-C**) and mEPSCs (**Fig EV5D-F**) between the wt, untreated and treated zQ175DN MSN groups.

These data indicate that exogenous cholesterol restores synaptic function in zQ175DN MSNs and that behavioral improvement correlates with normalization of electrophysiological parameters.

### Absence of long-term adverse effects of the treatment regimen

We also assessed the risk of adverse effects in the “2-cycle treatment”. For this purpose, a multiplex immunoassay (Bio-Plex) was used to simultaneously measure the level of 23 inflammatory cytokines in eight tissues (striatum, cortex, cerebellum, lung, liver, kidney, heart, spleen) and in plasma. We found an overall similar inflammatory profile among the three experimental groups with slight variations for only 2 pro-inflammatory analytes (eotaxin and creatine kinase, KC) in three tissues (striatum, cerebellum, and liver), suggesting that the treatment does not induce a relevant immune response (**Table EV4**).

Clinical observation of mice exposed to the different therapeutic regimens also revealed no fatalities in the groups nor did routine hematology reveal differences between the groups (**Table EV5**). Platelet count was higher in the zQ175DN+chol group than in the wt (*p* = 0.028) and zQ175DN (*p* = 0.011) groups. Large clumps of platelets were also detected on smears from some of the mice in the latter groups (wt, 2/6 mice; zQ175DN, 4/6 mice). A statistically significant difference in albumin concentration was detected between the zQ175DN+chol mice and wt mice (p = 0.019), and between zQ175DN+chol mice and zQ175DN mice (p = 0.025) (**Table EV6**). Despite a trend to increased values of ALT and AST, due to a single mouse with high values of both the enzymes, no statistically significant differences were seen in the other parameters. Overall, these data indicate good *in vivo* tolerability of 2 cycles of hybrid-g7-NPs-chol as no major adverse findings emerged.

No statistically significant differences in the total body weight and organs weight were found between zQ175DN and zQ175DN+chol groups (**Table EV7**). No major histopathological findings were found in any of the organs examined (**Table EV8**) and those occasionally present were interpreted as background or incidental lesions with no differences in their frequency or severity between groups.

In all cholesterol treated mice from the “2-cycle treatment”, numerous (5-6 per mouse), spherical, 2 to 3 mm in diameter, whitish, hard concretions were found loose in the abdominal cavity. No macroscopic lesions associated with the concretions were observed. Histological examination of these concretions revealed a central mineralized nucleus, intermingled with necrotic material, cholesterol clefts, variable number of foamy macrophages and occasional multinucleated giant cells surrounded by a fibrous capsule (foreign body granuloma) and covered externally by a monolayer of flattened mesothelium. These concretions were not or were rarely present in mice exposed to the other therapeutic regimens (in “early treatment” and “late treatment”) and their number and size depended on the timing and number of cycles (**Table EV9**).

## Discussion

There is now a substantial body of evidence indicating that delivering exogenous cholesterol to the HD brain or targeting its homeostasis can halt disease progression in mice. To date, cholesterol has been delivered to the brain of the rapidly progressing R6/2 mouse model via ip injection of cholesterol-loaded nanoparticles (Valenza *et al*, 2015a; Birolini *et al*, 2021b) or directly into the striatum via osmotic mini-pumps (Birolini *et al*, 2020) or even by nose-to-brain delivery of liposomes (Passoni *et al*, 2020). In a further strategy we used a gene therapy approach to overexpress the active form of SREBP2 in R6/2 striatal astrocytes *in vivo* (Birolini *et al*, 2021a), with the intention of promoting the transcription of genes in the cholesterol biosynthetic pathway in astrocytes, which produce most of the cholesterol present in the healthy adult brain and show reduced synthesis in the presence of mutant HTT (Valenza *et al*, 2015b). Others have targeted neuronal cholesterol catabolism by using an AAV vector overexpressing the neuronal-specific CYP46A1 gene in the striatum of R6/2 and zQ175 mice (carrying the neo cassette) (Kacher *et al*, 2019; Bousiccault *et al*, 2016). Activity of CYP46A1, which catalyzes the conversion of cholesterol into its 24S-OHC catabolite - which then passes the BBB and is detected in blood - is reduced in the brain of HD mice and in post-mortem brain tissues from HD patients (Valenza *et al*, 2007a; Valenza *et al*, 2010; Shankaran *et al*, 2017; Boussicault *et al*, 2016; Kacher *et al*, 2019). CYP46A1 gene therapy thus aims to increase cholesterol catabolism and reverse cholesterol accumulation of cholesterol - which was found increased in the striatum of R6/2 mice (Boussicault *et al*, 2016), but not in our studies on the same mice (Valenza *et al*, 2007a; Valenza *et al*, 2010; Shankaran *et al*, 2017; Birolini *et al*, 2020; Birolini *et al*, 2021a; Birolini *et al*, 2021b) nor in zQ175 mice (Kacher *et al*, 2019; this paper), - leading to a favorable outcome in HD mice. Thus, supplying the HD brain with exogenous cholesterol or enhancing its biosynthesis or catabolism led to improved neuronal function and lightening of phenotypes in two HD mouse models.

Here, we have proven the long-term beneficial effect of cholesterol administration through the most advanced formulation of brain-permeable NPs in counteracting cognitive decline, disease progression, motor defects, and neuropathological signs in the slowly progressing zQ175DN mouse model. To our knowledge this is the only study in which the therapeutic effect of a single molecule was confirmed in a long-term study (duration 10 months) in HD mice. Other studies examining long-term treatment effect include four HTT lowering studies and the aforementioned CYP46A1 gene therapy study (Kordasiewicz *et al*, 2012; Zeitler *et al*, 2019; Pikuleva *et al*, 2021; Marchionini *et al*, 2022; Kacher *et al*, 2019).

Importantly, we now demonstrate that cholesterol administration is equally effective in the presymptomatic and symptomatic stages. Comparing the “early treatment” with the “late treatment” or with the combination of both in the “2-cycle treatment” over nearly 1 year, we also conclude that treatment efficacy persists for 5 months after the last ip injection, regardless of the time of administration. This is in agreement with the slow release of cholesterol from the injected NPs which is completed in 5 months, as shown by Cy5-Bodipy imaging data. We also demonstrate that efficacy can be extended beyond 40 weeks (11 months) with the "2-cycle treatment”, therefore culminating in lasting recovery.

In this study, we used the zQ175DN mice backcrossed with inbred C57BL/6J mice and provided further characterization of the early stages of the disease in this mouse model. Previous data on these mice were obtained in a FVB background with the first signs of cognitive and motor deficits starting at 6-9 months (Southwell *et al*, 2016). Here we show that cognitive decline, neuromuscular (grip) defects, and functional impairment begin at 5 months (20 weeks) of age, a time that can be considered similar to the onset of a prodromal phase in humans (Rao *et al*, 2011; Paulsen *et al*, 2013; Ramos *et al*, 2017).

Cognitive impairment is often reported as the most debilitating aspect of HD that currently goes untreated. Our previous studies have shown that 15-20 µg of cholesterol, administered via g7-PLGA-NPs-chol (Valenza *et al*, 2015a) or with minipumps (Birolini *et al*, 2020), are sufficient to obtain complete prevention of cognitive decline and that higher doses of cholesterol maintain this benefit. In this study, we confirmed that the dose of cholesterol required to prevent or revert cognitive decline in HD mice is quite low, thus anticipating that a broad window of therapeutic intervention on cognition is available. Here we show that the attenuation of cognitive decline as measured by the NOR test in zQ175DN mice occurs with both “early treatment” and “late treatment” and is long-lasting with the “2-cycle treatment”. Instead, associative memory, as measured by the FC test, can only be rescued by the “2-cycle treatment”.

Recognition memory and associative memory in mice recruit both the hippocampus and surrounding regions of the cortex, but recognition memory is primarily associated with the cortico-striatal pathway (Milnerwood & Raymond, 2010; Darvas & Palmiter, 2009; Poulter *et al*, 2020). In contrast, the amygdala is mainly involved in FC-dependent associative memory (Krabbe *et al*, 2017). The stronger effect observed in NOR versus FC performance suggests that exogenous cholesterol acts primarily on cortico-striatal and hippocampal-striatal interactions (Goodroe *et al*, 2018). Thus, strategies aimed at increasing cholesterol biosynthesis may act as modifiers of cognitive symptoms that appear in the early phases of the disease. Importantly, Fig 3D shows that in all cholesterol-raising trials we performed to date, in a total of 381 mice, regardless of treatment, animal model, and time of delivery, the administration of exogenous cholesterol or improvement of its biosynthesis in the HD mouse brain completely normalizes cognition (the two AAV-CYP46AI mouse studies were not included as cognitive parameters were not measured, Kacher *et al*, 2019; Boussicault *et al*, 2016). We are not aware of any HD pre-clinical trial whose success has been documented in so many mice from two different models (i.e., a total of 127 treated and 130 untreated HD mice versus 124 controls were tested for cognitive parameters) and supported by five different strategies (i.e., first and second generation NPs, minipumps, AAV-SREBP2 and nose-to-brain delivery). The robustness of “brain-cholesterol” therapy in HD is bolstered by the demonstration that the number of muHTT aggregates, one of the key features of HD pathology, is also consistently reduced in animals given any of the cholesterol-raising strategies we have used thus far, with maximal effect observed in zQ175DN mice after “2-cycle treatment” with hybrid-g7-NPs-chol and no effect, as expected, when the same NPs are delivered to R6/2 mice due to the slow release of cholesterol from these NPs and fast disease progression of this mouse model.

Regarding clasping behavior - a measure of motor defect - recovery occurs only at 29 weeks of age and, at later times, only with the “2-cycle treatment”, while at the 20 weeks the cognitive defect was already healed. This is consistent with our previous data showing that motor recovery requires the release of a higher dose of cholesterol (Birolini *et al*, 2020). Consequently, complete recovery in the AC and RR tests coincided with the complete release of cholesterol from the NPs, while normalization in PC was maximal only when release was at its highest values (at 29 weeks of age).

Our data also indicate that exogenous cholesterol acts at the pre-synaptic level. In fact, its administration normalizes the frequency of sEPSCs, which reflects the number of synaptic contacts or excitability of pre-synaptic neurons, and of mEPSCs, which reflects the probability of stochastic release of synaptic vesicles from the presynaptic terminals and the number of synapses/active zones. These results support the idea that efficacy of exogenous cholesterol also depends on its ability to normalize synaptic defects in HD. A deeper understanding of the cell types and specific circuits associated with behavioral recovery in treated mice will help increase understanding of the biology of cholesterol-based recovery.

Extensive mass spectrometry-based studies demonstrated that cholesterol biosynthesis is impaired early in the HD brain and across different rodent models (Valenza *et al*, 2007a; Valenza *et al*, 2007b; Kacher *et al*, 2019; Shankaran *et al*, 2016; Valenza *et al*, 2010; Bossicault *et al*, 2016). In this study we confirmed a decrease in the steady-state level of cholesterol in the striatum of zQ175DN mice. We also show that cholesterol precursors and its major catabolite, 24S-OHC, are similarly reduced in the striatum and cortex of zQ175DN mice compared to wt mice. In our previous studies, a high dose of cholesterol infused directly into the striatum of R6/2 mice enhanced the biosynthesis of endogenous cholesterol (Birolini *et al*, 2020), probably through its conversion into 24S-OHC in neurons. 24S-OHC may act as a signaling molecule of neuronal cholesterol status and, by binding to LXRs on astrocytes, stimulate cholesterol efflux from these cells and, in turn, cholesterol biosynthesis (Albidayeva *et al*, 2006). However, in the present study, we did not detect increased steady-state levels of cholesterol precursors and 24S-OHC. It should be noted that these measurements were only taken only at the end of treatment, i.e. 5 or 10 months after the last ip injection, probably a late time to detect a modulation in endogenous of cholesterol.

Our previous results in R6/2 mice showed that cholesterol infusion (Birolini *et al*, 2020) or enhancement of endogenous cholesterol biosynthesis (Birolini *et al* 2021a) in the striatum promotes clearance of muHTT aggregates. Here we confirm this result in zQ175DN mice. After the “2-cycle treatment”, muHTT aggregates are barely present in both the striatum and cortex, suggesting that more cycles produce a greater and longer beneficial effect, and that exogenous cholesterol likely acts to increase muHTT clearance. The significant decrease of muHTT aggregates and the significant recovery of cognitive decline, disease progression and neuromuscular defects, suggest that clearance of muHTT aggregates after cholesterol administration may contribute to the animal’s behavioural improvement.

Finally, our analyses of inflammatory status and complete necropsy performed in HD mice after “2-cycle treatment” showed no significant changes or treatment-related lesions indicative of side effects. The intra-abdominal mineralized concretions observed in the zQ175DN+chol group are probably a foreign body reaction to the ip administration, which is the preferred route of systemic drug administration in mice but is less used in humans (Ramot *et al*, 2009; Alsina-Sanchis *et al*, 2021). Further studies on pharmacokinetics and regulatory toxicity with other routes of administration and greater translational significance, such as intravenous administration, are needed. Differences in platelet counts should be a preanalytical artifact induced by platelet clumping and were observed in all groups of animals. Similarly, the increased albumin concentration measured in the zQ175DN+chol group can be interpreted as a nonspecific finding, possibly due to moderate dehydration.

In summary, the results obtained in this study analyzing behavioral and neuropathological parameters on a total of 49 cholesterol-treated zQ175DN mice compared to 48 wt and 48 untreated zQ175DN mice, support the notion that cholesterol-based strategies act as modifiers of disease progression. We show that exogenous cholesterol is able to postpone symptoms when administered presymptomatically and to reverse symptoms when delivered in the symptomatic phase. We also show that 2 cycles of cholesterol administration resulted in a complete and sustained recovery of cognitive and motor deficits lasting almost a year. Furthermore, the slow release of exogenous cholesterol by NPs may mimic the low and constant daily cholesterol synthesis that occurs physiologically in the adult brain.

Given the robustness of the therapeutic impact of cholesterol delivery to the HD brain, the next challenge now is to optimize the strategies for its clinical application. A promising approach may come from the intranasal delivery of nanoparticles or liposomes, as proposed more recently (Passoni *et al*, 2020). Optimizing nose-to-brain delivery of cholesterol may help overcome the BBB and more easily deliver cholesterol to the brain. Furthermore, the new cholesterol-releasing products should have all the characteristics required by the market to cross the preclinical-clinical boundary and become a real therapy for HD patients.

## Materials and methods

### NPs production and characterization

Hybrid-g7-NPs loaded with Bodipy-chol and labelled with Cy5 (Cy5-hybrid-g7-NPs-Bodipy-chol), hybrid-g7-NPs loaded with d6-chol (hybrid-g7-NPs-d6-chol) and hybrid-g7-NPs-chol were prepared and characterized as previously described (Belletti *et al*, 2018; Birolini *et al*, 2021b). Briefly, all NPs were produced via a nanoprecipitation method by dissolving 25 mg of polymer and 25 mg of lipid in acetone and using a 0.5% w/V Pluronic® F68 water solution at 45°C as aqueous phase. For each batch, standard nanotechnological characterization was performed to assess NPs quality. Size (nm), polydispersity index (PDI; a value to estimate the uniformity in size of the NPs) and z-potential (mV; a measure reflecting the surface charge of the NPs) were measured with Photon Correlation Spectroscopy (Malvern Zetasizer ZS, Malvern, UK) at a concentration of 0.01 mg/mL (automatic laser attenuator, refractive index 1.59). The mean (± SD) values of these features of all NPs used in this study are herein summarized: size (nm): 249±38; PDI: 0.29±0.05; z-potential (mV): −30±7. The content of cholesterol in each batch was quantified via RP-HPLC-UV (Jasco, Cremella, Italy) using a system equipped with a Syncronis C18 column (250 × 4.6 mm; porosity 5 μm; Thermo Fisher Scientific, Waltham, MA, USA), performing a isocratic separation with 50:50 Acetonitrile:Ethanol as mobile phase (flow rate 1.2 mL/min, UV detector set at 210 nm). Cholesterol was extracted from lyophilized NPs before injection and PLGA was removed. The mean (±SD) cholesterol content was 28.71±3.498 mg of cholesterol for 100 mg of NPs. Data presented were calculated as mean of at least three independent measurements.

### zQ175DN colony management and treatment

zQ175DN mice on a C57BL6/J background were acquired by The Jackson Laboratories (B6.129S1-*Htt^tm1.1Mfc^*/190ChdiJ; Jax stock #029928; RRID:IMSR JAK:029928) and maintained by mating 6-weeks old male and female heterozygous mice with C57BL6/J mice (purchased from Charles River). The colony used in this study carried around 190 CAG repeats. Mice were weaned and then genotyped at 3 weeks of age (+/− 3 days). Mice were housed under standard conditions in enriched cage (22 ± 1°C, 60% relative humidity, 12 hours light/dark schedule, 3–4 mice/cage, with food and water ad libitum). After PCR genotyping, male and female mice were included and randomly divided into experimental groups. Littermates were included as controls. All animal experiments were approved and carried out in accordance with Italian Governing Law (D.lgs 26/2014; Authorization *n.581/2019-PR* issued July 29, 2019 by Ministry of Health and Authorization *n.714/2020-PR* issued July 21, 2020 by Ministry of Health); the NIH Guide for the Care and Use of Laboratory Animals (2011 edition) and EU directives and guidelines (EEC Council Directive 2010/63/UE).

7-weeks-old wild-type (wt) littermate mice (for cholesterol release and d6-chol quantification studies) and zQ175DN mice (for the behavioral experiments) were treated with 0.12 mg NPs/gr body weight (about 1.7 mg NPs/mouse; carrying about 493 µg of cholesterol) for ip injection. As controls, wt and zQ175DN littermates were treated with saline solution. In all trials, following ARRIVE guidelines, mice were assigned randomly, and sex was balanced between the various experimental groups; the animals of the same litter were divided in different experimental groups; investigator blinding was applied to in vivo procedures and all data collection.

### Bodipy analysis

Analysis was performed as described in (Birolini *et al*, 2021b). Briefly, 24 hours, 2 weeks, 10 weeks or 20 weeks after a single ip injection of Cy5-hybrid-g7-NPs-Bodipy-chol, mice were deeply anesthetized by ip injection of Avertin 2.5% (250 mg/kg body weight) and transcardially perfused with saline solution followed by PFA 4%. Brains were post-fixed for 2h in the same solution at 4°C (no longer to avoid fluorescence bleaching) and shifted into 30% sucrose to prevent ice crystal damage during freezing in OCT. 15 μm-thick brain coronal sections were counterstained with the nuclear dye Hoechst 33258 (1:10.000, Invitrogen) and then mounted under cover slips using Vectashield mounting medium (Vector Laboratories). The fluorescence signals of Cy5 and Bodipy-chol were acquired the following day. To quantify the released Bodipy-chol from hybrid-g7-NPs-chol, Volocity software was used using the plug-in “find objects” and “calculate object correlation” (*N* = 4 mice/time point; *N* = 6 images/mouse).

### Liquid chromatography-mass spectrometry (LC-MS) analysis for d6-chol quantification

The previously published validated method by Passoni et al. was used (Passoni *et al*, 2020). Briefly, an aliquot of plasma (50 μL) was diluted with 200 μL of ethanol containing 200 ng of beta-sitosterol, used as internal standard. Samples were vortexed and centrifuged at 13200 rpm for 15 min and 2 µL of the supernatants were directly injected into the LC-MS system. Each brain area and peripheral tissues were homogenized in 1 mL of PBS, containing 500 ng of internal standard. 200 µL of homogenate were extracted with 800 µL of ethanol, vortexed, and centrifuged for 15 min at 13200 rpm at 4 °C. 4 µL of each sample were injected into the LC-MS system.

D6-chol levels were determined using a 1200 Series HPLC system (Agilent Technologies, Santa Clara, CA, U.S.A.) interfaced to an API 5500 triple quadrupole mass spectrometer (Sciex, Thornhill, Ontario, Canada). The mass spectrometer was equipped with an atmospheric pressure chemical ionization (APCI) source operating in positive ion and multiple reaction monitoring (MRM) mode to measure the product ions obtained in a collision cell from the protonated [M – H2O]+ ions of the analytes. The transitions identified during the optimization of the method were m/z 375.3–152.1 (quantification transition) and m/z 375.3–167.1 (qualification transition) for D6-chol; m/z 397.3–147.1 (quantification transition) and m/z 397.3–161.1 (qualification transition) for β-sitosterol (IS). D6-chol and beta-sitosterol were separated on a Gemini C18 column (50 × 2 mm; 5 μm particle size), using an isocratic gradient in 100% methanol at 35 °C.

### Behavioral assessments

Novel Object Recognition (NOR) test, Paw Clasping (PC) test, Grip Strength (GS) test, and Activity Cage (AC) test were performed at 20, 29, 35, and 45 weeks of age for “early treatment” and “2-cycle treatment”, and at 40 and 49 weeks of age for “late treatment”. Rotarod (RR) test was performed at 46 weeks of age for “early treatment” and “2-cycle treatment” and at 50 weeks of age for “late treatment”. All these behavioral tests were conducted as described in (Birolini *et al*, 2021a).

Fear Conditioning (FC) test was performed at 48 weeks of age for “early treatment” and “2-cycle treatment” and at 51 weeks of age for “late treatment”. Contextual and cued fear conditioning were performed using the Fear Conditioning System (Ugo Basile, series 46000) according to the following protocol: on day 1, mice were acclimatized in the behavioral room 30 minutes before the start of the test. Mice were then exposed to a conditioned stimulus (CS) consisting in a 10-second 3.5 kHz tone delivered 150 seconds after the beginning of the test. The unconditioned stimulus (US) was a 2-second 0.5 mA foot shock that terminated together with the CS. The conditioning was repeated 3 times, every 150 seconds. After the last conditioning, animals were returned to their own cage. On day 2, contextual memory was assessed by placing mice in the same conditioning chamber for 5 minutes. The freezing response was evaluated by measuring the number of freezing, total time of freezing and latency to freeze. On day 3, cued fear conditioning was assessed using a modified conditioning chamber obtained by pasting patterned contexts onto the walls and floor. Once mice were placed in this modified chamber, the same protocol as of day 1 was applied, except for the electrical foot shock that was not administered. Number of freezing, time of freezing and latency to freeze were recorded. In the cued paradigm, the recording started 150 seconds after the beginning of the test, as the US was delivered at this point.

Y-maze test (YM) was performed at 30 and 48 weeks of age only for “2-cycle treatment”. A Y-shaped maze with three arms at a 120° angle from each other was used. After placing the mice in the center of the maze, they had free access to all three arms. If the mice chose an arm other than the one they came from, this choice was called an alternation, which was considered the correct response because mice prefer to investigate a new arm of the maze rather than returning to one that was previously visited. Indeed, returning to the previous arm was considered an error. The total number of arm entries and the sequence of entries were recorded in order to calculate the percentage of alternations according to the following formula: number of alternations / number of possible triads x 100. For a summary of animals used in the study and the tests performed, see **Table EV2**.

### muHTT aggregates analysis

zQ175DN mice from “early treatment” were sacrificed at 29 and 50 weeks of age and mice “late treatment” and “2-cycle treatment” were sacrificed at 29 and 50-51 weeks of age, respectively and the nalysis was performed as described in (Birolini *et al*, 2020). Mice (*N* = 3/group) were anesthetized by ip injection of Avertin 2.5% (250 mg/kg body weight) and transcardially perfused with saline solution followed by PFA 4%. Brains were post-fixed overnight in the same solution at 4°C and then in sucrose 30% to prevent ice crystal damage during freezing in OCT. Immunohistochemistry was performed on 15 μm coronal sections.

Epitopes were demasked at 98°C with NaCitrate (10 mM, pH 6) for 15 minutes. Tissue slices were then blocked and permeabilized with Normal Goat Serum 5% and Triton 0,5% for 1h at RT, followed by incubation with anti-EM48 primary antibody (1:100; Merck Millipore, cat n° MAB5374) for 3h at RT. Slices were washed 3 times (10 minutes each) with PBS + Triton 0.1% at RT and exposed to anti-mouse Alexa Fluor 488-conjugated goat secondary antibodies (1:500; Invitrogen) for 1h at RT. Sections were counterstained with the nuclear dye Hoechst 33258 (1:10.000, Invitrogen), washed 3 times (10 minutes each) with PBS + Triton 0.1% at RT and mounted under cover slips using Vectashield mounting medium (Vector Laboratories).

To count muHTT aggregates, confocal images were acquired with a LEICA SP5 laser scanning confocal microscope. Laser intensity and detector gain were maintained constant for all images and 5 to 10-z steps images were acquired at 40x. To count aggregates, 27 images/mice from 9 sections throughout the striatum and cortex were taken. To quantify the number and the size of aggregates, ImageJ software was used to measure the fluorescence. Images were divided into three-color channels and the same global threshold was set. “Analyze Particles” plugin (ImageJ) was used to count the number and the size of aggregates.

### Isotope dilution-gas chromatography-mass spectrometry (ID-GC-MS) analysis for neutral sterols and 24S-OHC in brain tissues

Animals (*N* = 4/group) were perfused with saline before isolating the striatum and cortex to avoid blood contamination. Brain tissues were frozen and kept at −80°C until their use. Frozen tissues were rapidly weighted and homogenized in ice with sterile PBS (400 µl for the striatum; 600 µl for the cortex) with an ultra-turrax homogenizer. In a screw-capped vial sealed with a Teflon-lined septum, 50 µl of tissue homogenates were mixed together with 500 ng of D4-lathosterol (CDN Isotopes), 500 ng of D6-desmosterol (Avantipolar Lipids), 100 ng of D6-lanosterol (Avantipolar Lipids), 400 ng of D7-24S-hydroxycholesterol (D7-24S-OHC) (Avantipolar Lipids) and 100 µg of epicoprostanol (Sigma-Merck) as internal standards, 50 µl of butylated hydroxytoluene (BHT) (5 g/L) and 25 µl of EDTA (10 g/l). Alkaline hydrolysis was allowed to proceed at room temperature (22°C) for 1 h in the presence of 1 M ethanolic potassium hydroxide solution under magnetic stirring. After hydrolysis, the neutral sterols (cholesterol, lathosterol, desmosterol and lanosterol) and 24S-OHC were extracted twice with 5 ml of hexane. The organic solvents were evaporated under a gentle stream of nitrogen and converted into trimethylsilyl ethers with BSTFA-1% TMCS (Cerilliant) at 70°C for 60 min. Analysis was performed by GC–MS on a Clarus 600 gas chromatograph (Perkin Elmer) equipped with Elite-5MS capillary column (30 m, 0.32 mm, 0.25 µm. Perkin Elmer) connected to Clarus 600C mass spectrometer (Perkin Elmer). The oven temperature program was as follows: the initial temperature 180°C was held for 1 min, followed by a linear ramp of 20°C/min to 270°C and then a linear ramp of 5°C/ min to 290°C, which was held for 10 min. Helium was used as carrier gas at a flow rate of 1 ml/min and 1 µl of sample was injected in splitless mode. Mass spectrometric data were acquired in selected ion monitoring mode. Peak integration was performed manually. Sterols and 24S-OHC were quantified against internal standards, using standard curves for the listed sterols. The weight of each sample was used to normalize raw data.

### Electrophysiological analysis

Experiments were performed on submerged brain slices obtained from mice from the “2-cycle treatment” (*N* = 3-4/group; 1-2 cells were recorded for each animal) at 53-55 weeks of age. Animals were anesthetized with isoflurane and transcardially perfused with ice-cold (<4°C), carboxygenated (95% O2 – 5% CO2) cutting solution (70 mM sucrose, 80 mM NaCl, 2.5 mM KCl, 26 mM NaHCO3, 15 mM glucose, 7 mM MgCl2, 1 mM CaCl2, 1.25 mM NaH2PO4). The brain was rapidly removed, and coronal slices (300 µm-thick) were cut at striatal level using a vibratome (DTK-1000, Dosaka EM, Kyoto, Japan). Slices were transferred into an incubating chamber filled with oxygenated ACSF (NaCl 125 mM, KCl 2.5 mM, NaHCO3 26 mM, Glucose 15 mM, MgCl2 1.3 mM, CaCl2 2.3 mM and NaH2PO4 1.25 mM) and allowed to equilibrate for 1 hour (30 minutes at 37°C and 30 minutes at room temperature). Slice were then transferred, one by one, to a submerged-style recording chamber for the whole-cell patch-clamp recordings. The chamber was mounted on an E600FN microscope (Nikon) equipped with 4× and 40× water immersion objectives (Nikon) and connected to a near-infrared CCD camera for cells visualization.

Data were obtained from MSNs (6 cells from 3 wt mice; 8 cells from 4 zQ175DN mice; 5 cells from 3 zQ175DN+chol mice), identified by their basic membrane properties (membrane capacitance, input resistance, and membrane resting potential) and firing pattern (delayed regular firing). The patch pipettes were produced from borosilicate glass capillary tubes (Hilgenberg GmbH) by using a horizontal puller (P-97, Sutter instruments) and filled with an intracellular solution containing K-gluconate 130 mM, NaCl 4 mM, MgCl_2_ 2 mM, EGTA 1 mM, creatine phosphate 5 mM, Hepes 10 mM, Na_2_ATP 2 mM, Na_3_GTP 0.3 mM (pH adjusted to 7.3 with KOH). The signals were amplified with a MultiClamp 700B amplifier (Molecular Devices) and digitized with a Digidata 1322 computer interface (Digitata, Axon Instruments Molecular Devices, Sunnyvale, CA). Data were acquired using the software Clampex 9.2 (Molecular Devices, Palo Alto, CA, U.S.A.), sampled at 20 kHz, and filtered at 2-10 kHz. The analysis of the membrane capacitance (Cm) and the input resistance (Rin) was performed using Clampfit 10.2 (Molecular Devices, Palo Alto, CA, U.S.A.). Cm was estimated from the capacitive current evoked by a −10 mV pulse, whereas Rin was calculated from the linear portion of the I-V relationship obtained by measuring steady-state voltage responses to hyperpolarizing and depolarizing current steps. The spontaneous excitatory postsynaptic currents (sEPSCs) – mediated by the activation of AMPA receptors as confirmed by their abolition following application of NBQX 10 µM – were recorded at a holding potential of −70 mV. Miniature EPSCs (mEPSCs) were derived in the presence of 1 μM TTX (Sigma-Aldrich). The off-line detection of the events was performed manually by using a custom-made software in Labview (National Instruments, Austin, TX, U.S.A.). The amplitudes of the events obeyed a lognormal distribution. Accordingly, the mean amplitude was computed as the peak of the lognormal function used to fit the distribution. Inter-event intervals (measured as time between two consecutive events) were distributed exponentially and the mean interval was computed as the tau value of the mono-exponential function that best fitted this distribution. The kinetic analysis of currents was performed by measuring rise time (time that current takes to activate from 10 to 90%) and time constant of decay (τd – time that current takes to deactivate exponentially to 37% of its peak value).

### Bio-Plex analysis

To measure cytokines and chemokines, mice from the “2-cycle treatment” (*N* = 4/group) were anesthetized by ip injection of Avertin 2.5% (250 mg/kg body weight). Blood was collected from the retro-orbital sinus in a tube containing 20 µL of heparin 2% and centrifuged at 4000 rpm in a bench-top centrifuge for 15 minutes to collect the plasma. Tissues (striatum, cortex, cerebellum, lung, liver, kidney, heart, and spleen) were isolated and frozen. 5 mg of each tissue were homogenized using a tissue grinder in 0.5 mL of lysing solution composed of 20 µL of factor 1, 10 µL of factor 2, 4995 µL of cell lysis buffer and 20 µL of PMSF 500 mM (Bio-Plex^®^ Cell Lysis Kit, Biorad, #171304011). The lysate was frozen at −80°C for 2 minutes and then, following thawing on ice, it was sonicated at 40% for 20 seconds and centrifuged at 4500 rcf at 4°C for 4 minutes to collect the supernatant.

The supernatant was quantified using DC™ Protein Assay Kit I (Biorad, #5000111) and samples were diluted to a final concentration of 500 µg/mL. To perform the Bio-Plex assay, 150 µL of assay buffer were added to 150 µL of samples. Concerning the plasma, samples were centrifuged at 1500 rcf at 4°C for 5 minutes. 60 µL of assay buffer and 120 µL of sample diluent were added to 60 µL of plasma.

Cytokine levels were measured by using a Bio-Plex murine cytokine 23-Plex assay kit (Biorad, #M60009RDPD) which evaluated the levels of: Eotaxin, IL-1a, IL-1b, IL-2, IL-3, IL-4, IL-5, IL-6, IL-9, IL-10, IL-12(p40), IL-12(p70), IL-13, IL-17, TNF-alpha, granulocyte colony-stimulating factor (G-CSF), granulocyte/macrophage colony-stimulating factor (GM-CSF), IFN-gamma, KC, RANTES, macrophage inflammatory protein (MIP-1a and MIP-1b) and monocyte chemotactic protein-1.

Briefly, magnetic beads were added to a 96-well plate which was fixed on a handheld magnetic washer. Samples were added and the plate was shacked at 300 rpm for 30 minutes at room temperature. After 3 washing steps, detection antibodies were added, and the plate was shacked at 300 rpm for 30 minutes at room temperature. After 3 washing steps, streptavidin-PE was added, and the plate was shacked at 300 rpm for 10 minutes at room temperature. After 3 washing steps, assay buffer was added and the cytokines levels were read on the Luminex 200 System, Multiplex Bio-Assay Analyzer and quantified based on standard curves for each cytokine in the concentration range of 1–32,000 pg/mL. The analytes concentrations specified for the eight-point standard dilution set included in the kit have been selected for optimized curve fitting using the five-parameter logistic regression in Bio-Plex ManagerTM software. These curves were used to interpolate the measured concentration. Data were normalized over the mean of the wt group.

### *In vivo* evaluation of safety: hematology, clinical chemistry, and histopathology

17 mice from “2-cycle treatment” (*N* = 5-6/group) underwent pathological evaluation to exclude adverse effects of the treatment. Mice were sacrificed at 50 weeks of age. Peripheral blood was sampled from the retro-orbital sinus and collected in Eppendorf Tubes® containing 20 µL of Heparin 2%. Blood samples were analyzed by means of a laser-based hematology analyzer (Sysmex XT-2000iV, Kobe, Japan). Blood smears were also prepared, stained with May Grunwald Giemsa and examined. Plasma obtained from blood samples was used to measure biochemical analytes by means of an automated spectrophotometer (BT3500, Biotecnica Instruments SPA, Rome, Italy). The following analytes were measured: cholesterol, triglycerides, glucose, total protein, albumin, creatinine, ALT and AST.

A complete necropsy was also performed. The total body weight was recorded, along with the weight of liver, spleen, kidneys, heart, and testes. The weight of the organs was normalized for the total body weight. Femoral bone marrow smears were also prepared, stained with May Grunwald Giemsa and examined. The following organs were collected and routinely processed for histopathology: brain, heart, lungs, liver, kidneys, spleen, salivary glands with mandibular lymph nodes, skin, skeletal muscle, testes, uterus, and ovaries. Hematoxylin and Eosin slides were examined by two pathologists blinded to the treatment groups.

### Heat Maps and Principal Component Analyses

Heat maps summarizing the behavior were created in excel using conditional formatting. For each set of data, a color scale was selected to define the best (light blue) or worst (dark blue) performance.

Principal component analysis (PCA) at 29 weeks of age and at 45-49 weeks of age was calculated using the R package stats (version 4.1.1) and visualized using factoextra (version 1.0.7), visualizing also the convex hull of the set of points belonging to each group of mice. At 29 weeks of age, PCA was performed using the following variables: Nor, Paw Clasping, Grip Strength, Activity Cage (global) and Activity Cage (distance). At 45-49 weeks of age, two more variables (Rotarod and Fear Conditioning) were considered.

### Meta-analysis and data-visualization (NOR test and muHTT aggregates)

All raw data of the Discrimination Index obtained with the NOR test performed in HD mice (R6/2 and zQ175DN) treated with different cholesterol-based strategies (Valenza *et al*, 2015a; Birolini *et al*, 2020; Birolini *et al*, 2021a; Birolini *et al*, 2021b and this study) were collected and re-analyzed. Data were visualized using a dotplot also showing their distribution, the differences between groups have been assessed using a pairwise t-test, multiple testing correction was performed using the Bonferroni method. The plot was generated using ggplot2 (version 3.4.0).

All raw data on the number of muHTT aggregates in the striatum of HD mice (R6/2 and zQ175DN) treated with different cholesterol-based strategies (Valenza *et al*, 2015a; Birolini *et al*, 2020; Birolini *et al*, 2021a; Birolini *et al*, 2021b and this study) were collected and the mean values for each group were normalized to 100 (corresponding to the mean of muHTT aggregates in untreated HD mice). Then, the normalized data were represented using a circular barplot to show the mean number of aggregates in the different treatments. The plot was generated using ggplot2 (version 3.4.0).

### Statistics

G*Power (https://www.psychologie.hhu.de/arbeitsgruppen/allgemeine-psychologie-und-arbeitspsychologie/gpower) was used to compute statistical power analysis in order to pre-determine group allocation, data collection, and all related analyses. Prism 9 (GraphPad software) was used to perform all statistical analyses. Grubbs’ test was applied to identify outliers. For each set of data to be compared, we determined whether data were normally distributed to select parametric or not parametric statistical tests. Data are presented as mean±standard error of the mean (mean±SEM). The limit of statistical significance was set at p-value < 0.05. The specific statistical test used is indicated in the legend of the figures.

## Data availability

This study does not include data deposited in public repositories. Data are available on request to the corresponding authors.

## Acknowledgements

The authors acknowledge Dr. Stefania Antonini, delegate of the University of Milan for the animal care, the designed veterinarians and the Charles River technical personnel of the animal facility located at L.I.T.A. (Segrate) for their support in the management of mouse colonies, and Prof. Flavia Antonucci and Dr. Clara Cambria for advice about YM test.

## Author contributions

MV, GB and EC conceived the study; GB managed the colonies and performed the treatments; GB and MV designed the treatment regimens, performed behavioral tests, sacrificed mice and collected tissues for all the subsequent analyses; IO, BR, JTD, MVa, RC, GT prepared and characterized all the NPs used in the study and contributed to the design of in vivo experiments; GB performed immunostaining experiments and provided confocal images and quantification; MRN performed the PCA analyses and metadata analysis of cognitive task and number of aggregates in different trials; FT, CCan, AT and GBi performed and analyzed the electrophysiological recordings; CC, FT and VL performed and analyzed mass spectrometry experiments; AP, MF, LC, RB and MS performed and analyzed mass spectrometry analysis for d6-chol quantification; ES, AC, LM performed histopathology and SP performed hematology and clinical chemistry; MV and GB collected study data and performed statistical analysis; MV and EC oversaw and coordinated responsibility for all research activities and their performance and provided experimental advice throughout the work. EC secured the fundings and the execution of the entire project. MV, GB and EC wrote the paper.

---

**Figure EV1.**
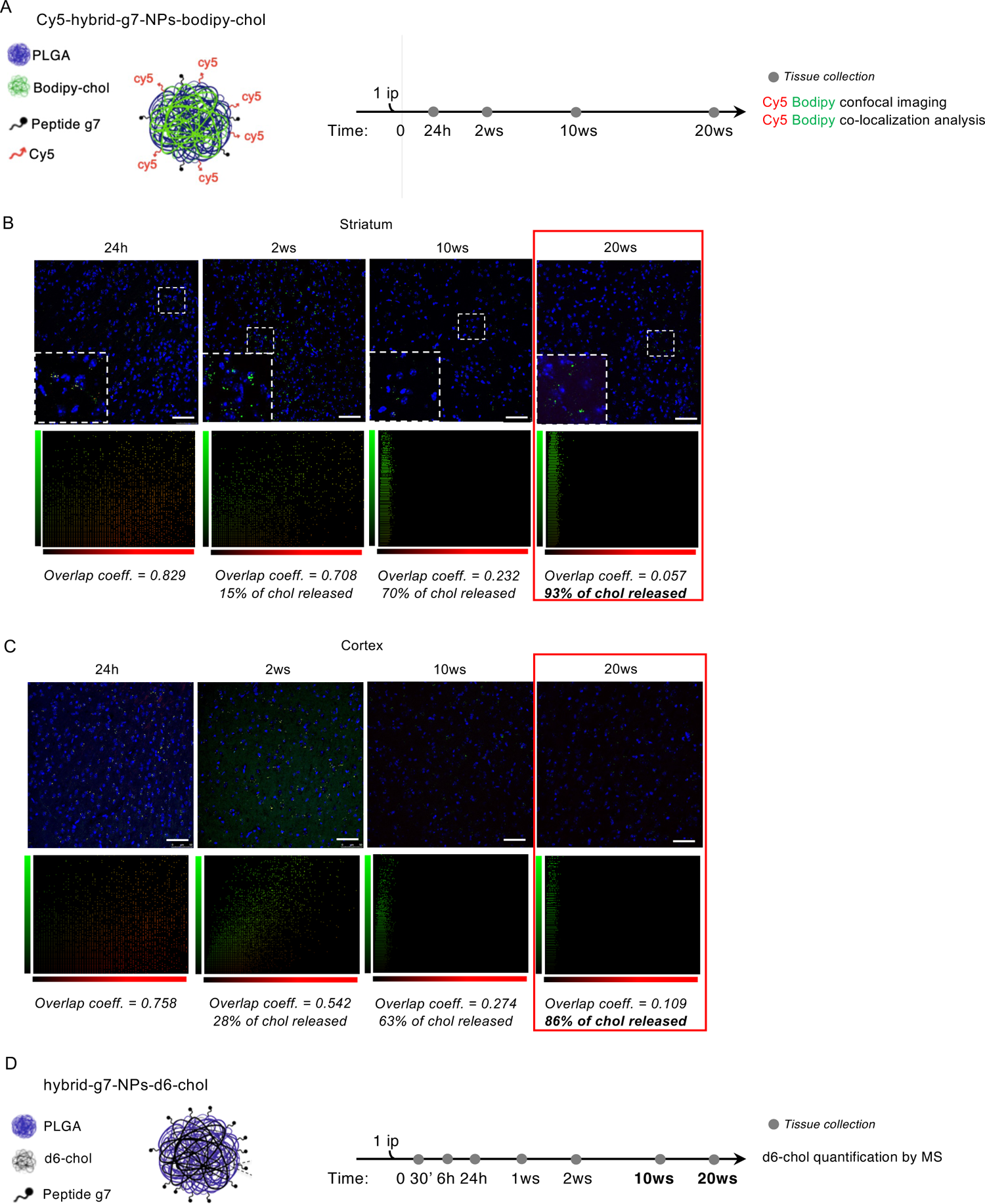
Cholesterol release and quantification by hybrid-g7-NPs-chol in wt mice over time. **A** Experimental paradigm: wt mice (*N* = 4/time point) were treated with a single ip injection of Cy5-hybrid-g7-NPs-bodipy-chol and sacrificed at different time points. Brains were collected for the analysis. **B–C** Representative confocal images of brain slices (striatum in B, cortex in C) from wt mice after ip injection of hybrid-Cy5-g7-NPs-bodipy-chol sacrificed after 24 h, 2 weeks, 10 weeks or 20 weeks and relative co-localization of bodipy-chol and g7-NPs. Hoechst was used to counterstain nuclei (Ho, blue). Scale bar is 50 μm. **D** Experimental paradigm: wt mice (*N* = 4/time point) were treated with a single ip injection of hybrid-g7-NPs-d6-chol and sacrificed at different time points. Different cerebral regions were collected for mass spectrometry analysis.

**Figure EV2.**
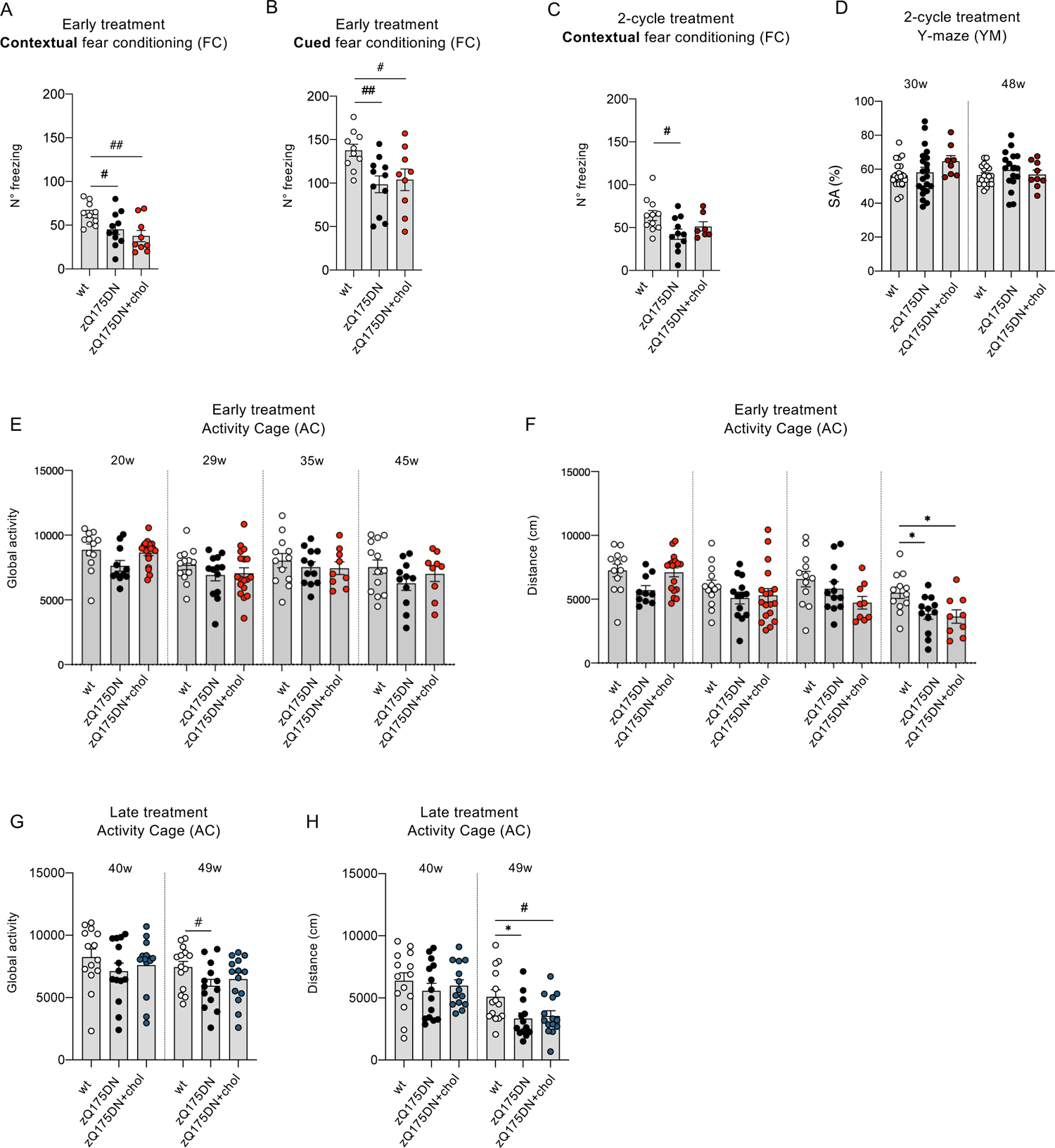
Cognitive and motor abilities of zQ175DN mice after the different treatments. **A-B** N° of freezing in the fear conditioning (FC) test in the contextual (A) and cued paradigm (B) from the “early treatment” at 48 weeks of age. **C** N° of freezing in the fear conditioning (FC) test in the contextual paradigm from the “2-cycle treatment” at 48 weeks of age. **D** Y-maze (YM) test from the “2-cycle treatment” at 30-48 weeks of age. Spontaneous alternation (SA) was calculated by dividing the n° of alternations by n° of possible triads x 100. **E–F** Global motor activity (E) and total distance traveled (in cm) (F) in an Activity Cage (AC) test from the “early treatment”. **G–H** Global motor activity (G) and total distance traveled (in cm) (H) in an Activity Cage (AC) test from the “late treatment”. Data information: data in A–H are shown as scatterplot graphs with mean±SEM. Each dot corresponds to the value obtained from each animal. Statistics: one-way ANOVA with Tuckey post-hoc test (*p<0.05); Student’s t-test (#p<0.05; ##p<0.01).

**Figure EV3.**
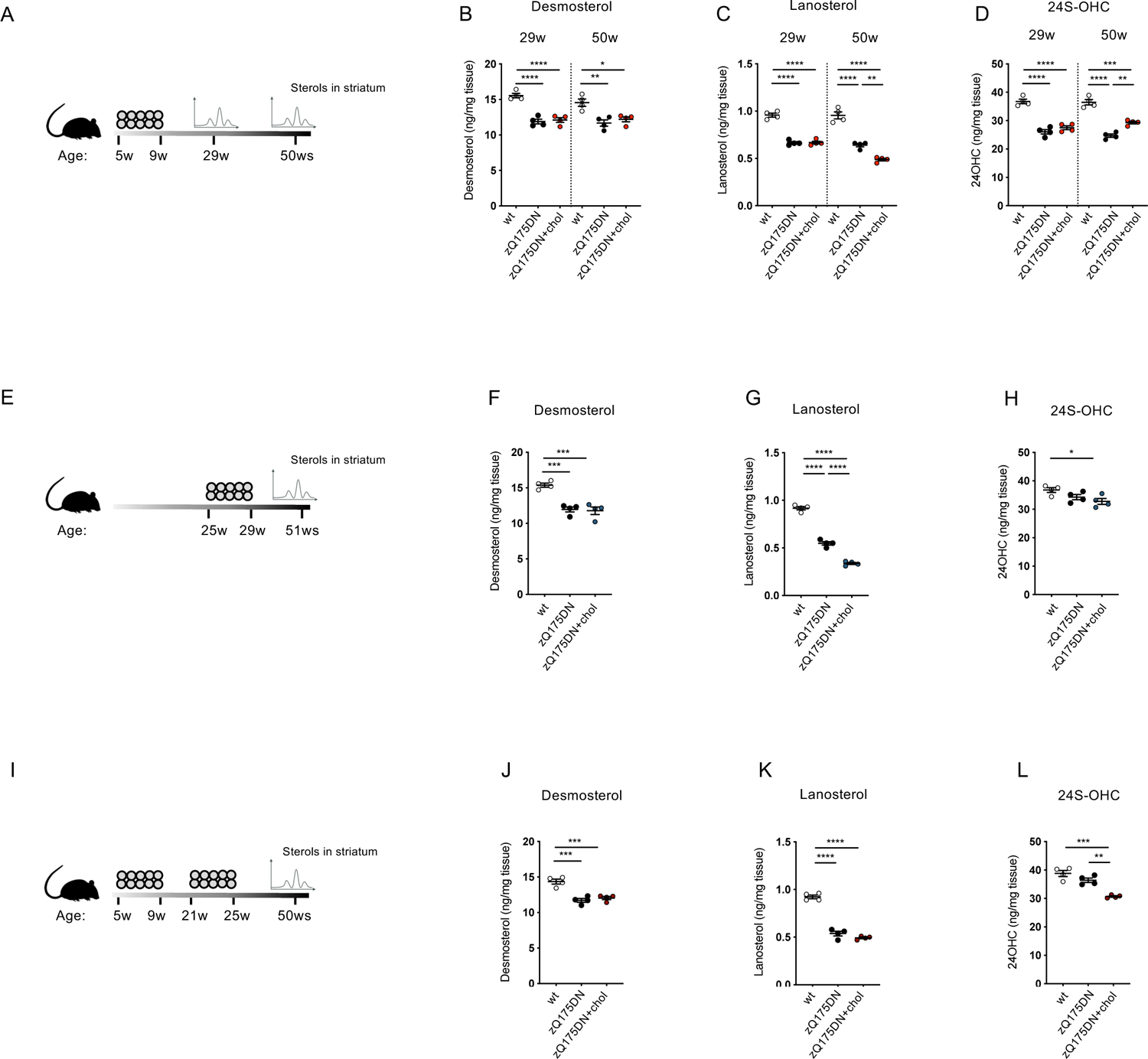
Sterols in the striatum of zQ175DN mice after the different treatments. **A-D** Mass spectrometry analysis performed in the striatum from the “early treatment” (*N* = 4 mice/group). Scheme of the treatment (A); level of desmosterol (B), lanosterol (C), and 24S-OHC (D) in the striatum at 29 and 50 weeks. **E-H** Mass spectrometry analysis performed in the striatum from the “late treatment” (*N* = 4 mice/group). Scheme of the treatment (E); level of desmosterol (F), lanosterol (G), and 24S-OHC (H) in the striatum at 51 weeks. **I-L** Mass spectrometry analysis performed in the striatum from the “2-cycle treatment” (*N* = 4 mice/group). Scheme of the treatment (I); levels of desmosterol (J), lanosterol (K), and 24S-OHC (L) in the striatum at 50 weeks. Data information: data in B–D, F–H, and J–L are shown as scatterplot graphs with mean±SEM. Each dot corresponds to the value obtained from each animal. Statistics: one-way ANOVA with Tuckey post-hoc test (*p<0.05; **p<0.01; ***p<0.001; ****p<0.0001).

**Figure EV4.**
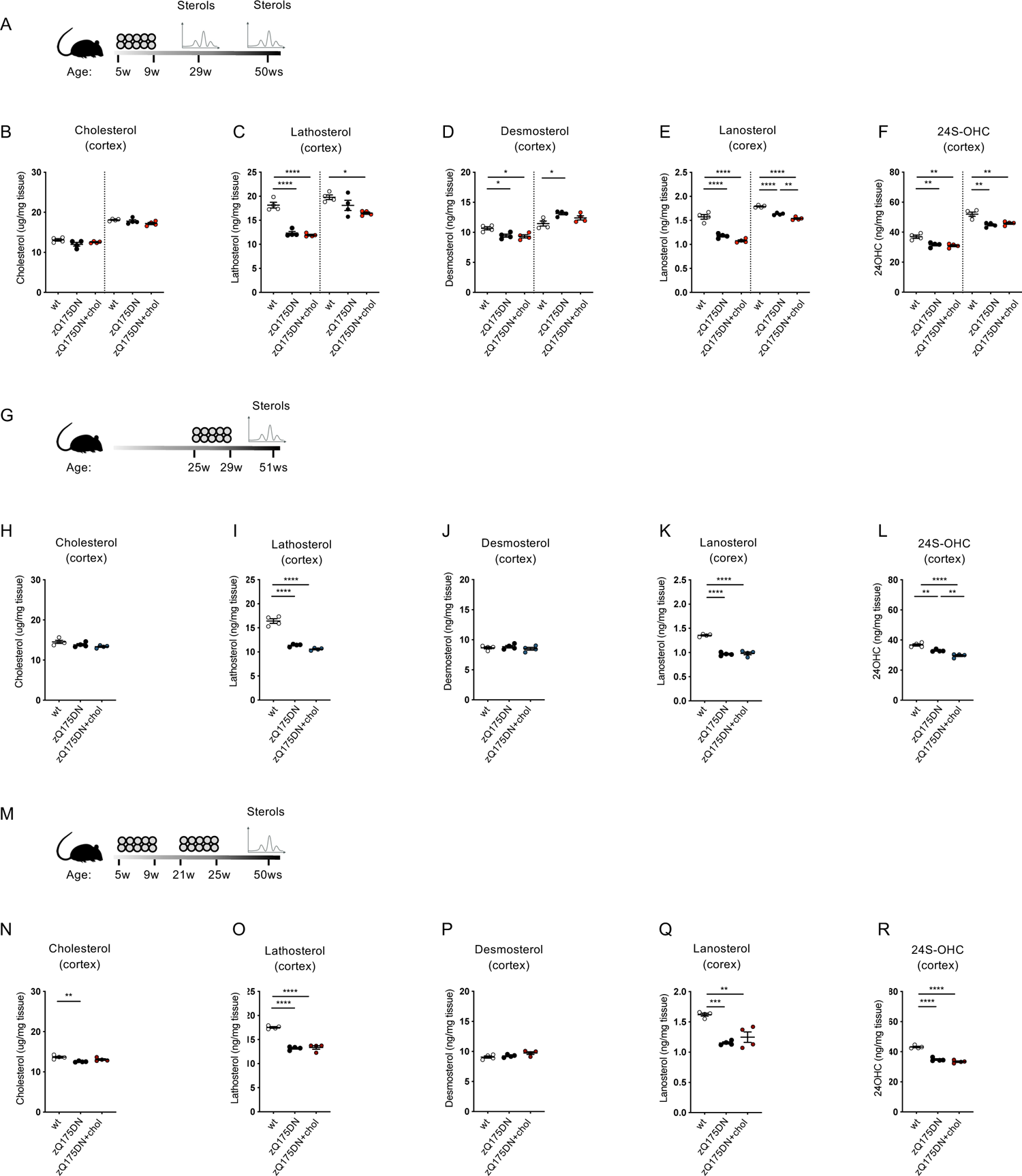
Sterols in the cortex of zQ175DN mice after the different treatments. **A-F** Mass spectrometry analysis performed in the cortex from the “early treatment” (*N* = 4 mice/group). Scheme of the treatment (A); level of cholesterol (B), lathosterol (C), desmosterol (D), lanosterol (E), and 24-hydrocycholesterol (F) in the cortex at 29 and 50 weeks. **G-L** Mass spectrometry analysis performed in the cortex from the “late treatment” (*N* = 4 mice/group). Scheme of the treatment (G); levels of cholesterol (H), lathosterol (I), desmosterol (J), lanosterol (K), and 24S-OHC (L) in the cortex at 51 weeks. **M-R** Mass spectrometry analysis performed in the cortex from the “2-cycle treatment” (*N* = 4 mice/group). Scheme of the treatment (M); level of cholesterol (N), lathosterol (O), desmosterol (P), lanosterol (Q), and 24S-OHC (R) in the cortex at 50 weeks. Data information: data in B–F, H–L, and N–R are shown as scatterplot graphs with mean±SEM. Each dot corresponds to the value obtained from each animal. Statistics: one-way ANOVA with Tuckey post-hoc test (*p<0.05; **p<0.01; ***p<0.001; ****p<0.0001).

**Figure EV5.**
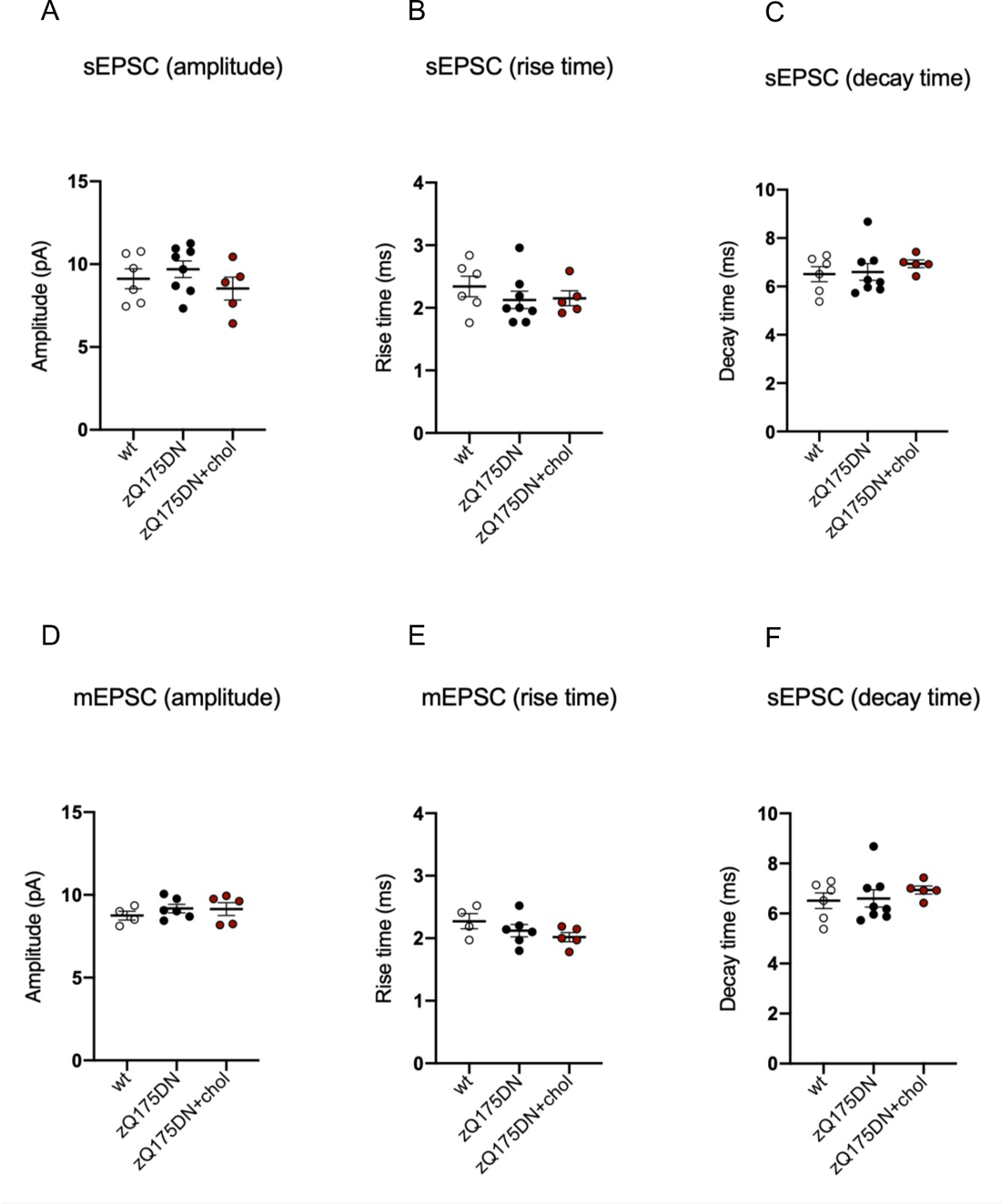
Electrophysiological features of MSNs from zQ175DN mice after the “2-cycle treatment”. **A–F** Average amplitude, rise time, and decay time of sEPSCs (A-C) and mEPSCs (D-F) recorded from MSNs of wt (6 cells from *N* = 3 mice), zQ175DN (8 cells from *N* = 4 mice) and zQ175DN+chol mice (5 cells from *N* = 3 mice). Data information: data in A-F are shown as scatterplot graphs with mean±SEM. Each dot corresponds to the value obtained from each cell.

**Table EV1:**
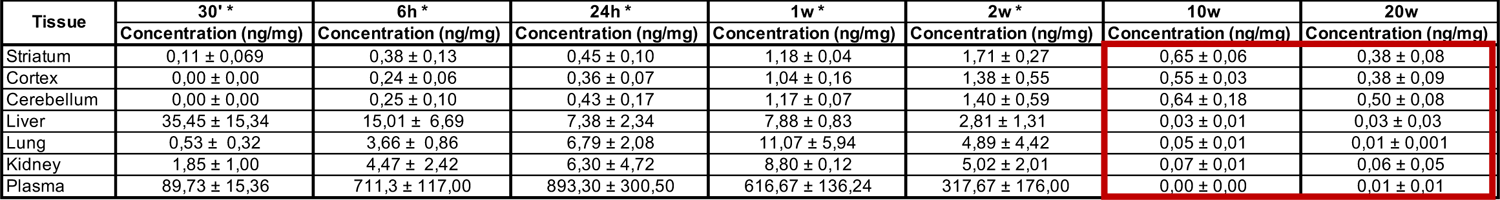
Level of d6-chol in striatum, cortex, cerebellum, liver, lung, kidney, and plasma measured by LC-MS (*N* = 3/group). Data information: data are expressed as mean±SEM. Red boxes highlight values measured at 10 and 20 weeks after ip injection in each tissue. Data indicated with * were previously published (Birolini et al., JCR 2021).

**Table EV2:** Number of animals used in each test performed in all the trials of this study.

|  | Early treatment |  |  |  |  |  |  |  |  |  |  |  | n. of tests performed (total) |
| --- | --- | --- | --- | --- | --- | --- | --- | --- | --- | --- | --- | --- | --- |
|  | wt |  |  |  | zQ175DN |  |  |  | zQ175DN+chol |  |  |  |  |
|  | 20w | 29w | 35w | 45-48w | 20w | 29w | 35w | 45-48w | 20w | 29w | 35w | 45-48w |  |
| Novel Object Recognition test (NOR) | 13 | 13 | 12 | 12 | 10 | 13 | 12 | 12 | 18 | 18 | 9 | 9 | 151 |
| Paw Clasping test (PC) | 13 | 12 | 12 | 12 | 11 | 13 | 12 | 12 | 18 | 18 | 9 | 9 | 151 |
| Grip Strength test (GC) | 13 | 13 | 12 | 12 | 11 | 13 | 12 | 12 | 18 | 18 | 9 | 9 | 152 |
| Activity Cage test (AC) | 13 | 13 | 12 | 12 | 10 | 13 | 12 | 12 | 18 | 18 | 9 | 9 | 151 |
| Rotarod test (RR) | - | - | - | 13 | - | - | - | 13 | - | - | - | 9 | 35 |
| Fear Conditioning test (FC) | - | - | - | 10 | - | - | - | 11 | - | - | - | 9 | 30 |

|  | Late treatment |  |  |  |  |  | n. of tests performed (total) |
| --- | --- | --- | --- | --- | --- | --- | --- |
|  | wt |  | zQ175DN |  | zQ175DN+chol |  |  |
|  | 40w | 49-51w | 40w | 49-51w | 40w | 49-51w |  |
| Novel Object Recognition test (NOR) | 14 | 14 | 14 | 13 | 14 | 14 | 83 |
| Paw Clasping test (PC) | 14 | 14 | 14 | 13 | 14 | 14 | 83 |
| Grip Strength test (GC) | 14 | 14 | 14 | 13 | 14 | 14 | 83 |
| Activity Cage test (AC) | 14 | 14 | 14 | 13 | 14 | 14 | 83 |
| Rotarod test (RR) | - | 14 | - | 13 | - | 14 | 41 |
| Fear Conditioning test (FC) | - | 13 | - | 13 | - | 13 | 39 |

|  | 2-cycle treatment |  |  |  |  |  |  |  |  |  |  |  | n. of tests performed (total) |
| --- | --- | --- | --- | --- | --- | --- | --- | --- | --- | --- | --- | --- | --- |
|  | wt |  |  |  | zQ175DN |  |  |  | zQ175DN+Chol |  |  |  |  |
|  | 20w | 29-30w | 35w | 45-48w | 20w | 29-30w | 35w | 45-48w | 20w | 29-30w | 35w | 45-48w |  |
| Novel Object Recognition test (NOR) | 15 | 18 | 19 | 18 | 13 | 17 | 18 | 18 | 17 | 17 | 17 | 17 | 204 |
| Paw Clasping test (PC) | 15 | 18 | 19 | 18 | 13 | 17 | 18 | 18 | 17 | 17 | 17 | 17 | 204 |
| Grip Strength test (GC) | 15 | 18 | 19 | 18 | 13 | 17 | 18 | 18 | 17 | 17 | 17 | 17 | 204 |
| Activity Cage test (AC) | 15 | 18 | 19 | 18 | 13 | 16 | 18 | 18 | 17 | 17 | 17 | 17 | 203 |
| Rotarod test (RR) | - | - | - | 17 | - | - | - | 13 | - | - | - | 17 | 47 |
| Fear Conditioning test (FC) | - | - | - | 11 | - | - | - | 11 | - | - | - | 7 | 29 |
| Y-maze test (YM) | - | 21 | - | 18 | - | 21 | - | 17 | - | 8 | - | 9 | 94 |

**Table EV3:** Quantification of the n° of muHTT aggregates in brain coronal slices of HD mice to measure the effect of the different strategies aimed at restoring cholesterol level/synthesis at the time points where a behavioural effect was observed. The mean values for each group were normalized to 100.

| R6/2_SREBP2 gene therapy |  |  |  | R6/2_minipumps |  |  |  | R6/2_NPs |  |  |
| --- | --- | --- | --- | --- | --- | --- | --- | --- | --- | --- |
| R6/2 + AAV-Tom<br>Contralateral<br>hemisphere | R6/2 + AAV-Tom<br>Infused<br>hemisphere | R6/2 + AAV-hBP2<br>Contralateral<br>hemisphere | R6/2 + AAV-hBP2<br>Infused<br>hemisphere | R6/2 + ACSF<br>Contralateral<br>hemisphere | R6/2 + ACSF<br>Infused<br>hemisphere | R6/2 + Chol<br>Contralateral<br>hemisphere | R6/2 + Chol<br>Infused<br>hemisphere | R6/2<br>+<br>saline | R6/2<br>+<br>g7-PLGA-NPs-chol | R6/2<br>+<br>g7-hybrid-NPs-chol |
| 104 | 122 | 71 | 88 | 38 | 94 | 97 | 23 | 142 | 97 | 52 |
| 120 | 85 | 119 | 96 | 108 | 33 | 56 | 18 | 75 | 138 | 112 |
| 118 | 71 | 90 | 94 | 73 | 54 | 56 | 32 | 157 | 64 | 106 |
| 50 | 87 | 81 | 82 | 105 | 49 | 56 | 41 | 106 | 151 | 130 |
| 62 | 85 | 73 | 84 | 41 | 39 | 76 | 23 | 90 | 68 | 132 |
| 58 | 74 | 77 | 58 | 80 | 54 | 56 | 51 | 81 | 64 | 46 |
| 133 | 84 | 136 | 147 | 26 | 33 | 38 | 25 | 129 | 70 | 147 |
| 169 | 49 | 127 | 121 | 11 | 25 | 14 | 32 | 141 | 54 | 100 |
| 85 | 139 | 127 | 155 | 153 | 36 | 351 | 29 | 81 | 109 | 66 |
| 139 | 86 | 134 | 123 | 48 | 50 | 243 | 17 | 133 | 76 | 84 |
| 116 | 96 | 82 | 119 | 157 | 121 | 32 | 23 | 140 | 95 | 123 |
| 85 | 97 | 137 | 123 | 270 | 31 | 8 | 11 | 83 | 74 | 54 |
| 123 | 102 | 82 | 79 | 90 | 113 | 123 | 1 | 62 | 117 | 49 |
| 110 | 64 | 91 | 94 | 242 | 112 | 352 | 2 | 83 | 79 | 118 |
| 97 | 66 | 65 | 65 | 264 | 81 | 188 | 2 | 74 | 66 | 134 |
| 75 | 99 | 100 | 84 | 52 | 117 | 39 | 0 | 78 | 88 | 104 |
| 56 | 95 | 102 | 90 | 75 | 125 | 277 | 0 | 89 | 133 | 101 |
| 56 | 87 | 108 | 89 | 68 | 124 | 45 | 2 | 90 | 76 | 73 |
| 70 | 95 | 108 | 69 | 106 | 45 | 68 | 12 | 70 | 86 | 108 |
| 83 | 67 | 125 | 92 | 113 | 28 | 44 | 44 | 84 | 85 | 95 |
| 113 | 75 | 99 | 42 | 33 | 47 | 57 | 44 | 131 | 82 | 122 |
| 86 | 68 | 80 | 32 | 48 | 158 | 64 | 14 | 117 | 91 | 109 |
| 92 | 121 | 97 | 59 | 152 | 197 | 92 | 17 | 72 | 108 | 131 |
| 119 | 102 | 81 | 71 | 72 | 118 | 41 | 35 | 140 | 113 | 81 |
| 136 | 111 | 99 | 47 | 77 | 97 | 55 | 14 | 67 | 110 | 97 |
| 145 | 114 | 119 | 34 |  | 19 | 75 | 13 | 104 | 121 | 115 |
|  | 141 | 94 | 28 |  | 165 |  | 22 | 80 | 111 | 130 |
|  |  | 117 | 75 |  |  |  |  |  |  |  |
|  |  | 89 | 79 |  |  |  |  |  |  |  |
|  |  | 150 | 42 |  |  |  |  |  |  |  |
|  |  | 60 | 55 |  |  |  |  |  |  |  |
|  |  | 103 | 78 |  |  |  |  |  |  |  |
|  |  | 72 | 8 |  |  |  |  |  |  |  |
|  |  | 106 | 27 |  |  |  |  |  |  |  |
|  |  | 121 | 49 |  |  |  |  |  |  |  |
|  |  | 83 | 51 |  |  |  |  |  |  |  |
|  |  | 112 | 47 |  |  |  |  |  |  |  |
|  |  | 85 | 63 |  |  |  |  |  |  |  |
|  |  | 68 | 44 |  |  |  |  |  |  |  |
|  |  | 114 | 60 |  |  |  |  |  |  |  |
|  |  | 106 | 72 |  |  |  |  |  |  |  |
|  |  | 120 | 83 |  |  |  |  |  |  |  |
|  |  | 98 | 55 |  |  |  |  |  |  |  |
|  |  | 94 | 55 |  |  |  |  |  |  |  |
|  |  | 103 |  |  |  |  |  |  |  |  |

| zQ175DN_g7-hybrid-NPs-chol |  |  |  |  |  |  |  |
| --- | --- | --- | --- | --- | --- | --- | --- |
| zQ175DN + saline<br>Early treatment<br>29w | zQ175DN + NPs<br>Early treatment<br>29w | zQ175DN + saline<br>Early treatment<br>50w | zQ175DN + NPs<br>Early treatment<br>50w | zQ175DN + saline<br>Late treatment<br>51w | zQ175DN + NPs<br>Late treatment<br>51w | zQ175DN + saline<br>2 cycles treatment<br>50w | zQ175DN + NPs<br>2 cycles treatment<br>50w |
| 85 | 58 | 39 | 81 | 69 | 5 | 128 | 1 |
| 73 | 85 | 62 | 181 | 94 | 35 | 104 | 5 |
| 94 | 74 | 113 | 167 | 88 | 75 | 74 | 4 |
| 117 | 104 | 85 | 73 | 67 | 62 | 116 | 6 |
| 98 | 92 | 75 | 95 | 79 | 19 | 94 | 4 |
| 104 | 98 | 86 | 142 | 98 | 83 | 83 | 6 |
| 100 | 86 | 118 | 133 | 77 | 49 | 83 | 7 |
| 119 | 76 | 84 | 137 | 100 | 132 | 141 | 3 |
| 88 | 82 | 98 | 149 | 82 | 88 | 73 | 2 |
| 134 | 79 | 97 | 51 | 123 | 48 | 120 | 6 |
| 163 | 98 | 113 | 58 | 80 | 51 | 213 | 5 |
| 92 | 71 | 128 | 118 | 80 | 123 | 51 | 3 |
| 88 | 55 | 150 | 115 | 104 | 37 | 149 | 5 |
| 143 | 84 | 99 | 63 | 113 | 36 | 86 | 12 |
| 97 | 56 | 88 | 56 | 54 | 135 | 96 | 4 |
| 131 | 84 | 107 | 60 | 91 | 86 | 67 | 5 |
| 138 | 77 | 117 | 50 | 94 | 64 | 126 | 3 |
| 68 | 147 | 71 | 59 | 97 | 62 | 60 | 7 |
| 68 | 69 | 160 | 167 | 117 | 67 | 98 | 2 |
| 63 | 58 | 81 | 72 | 147 | 97 | 104 | 2 |
| 38 | 101 | 114 | 111 | 104 | 38 | 73 | 3 |
| 103 | 38 | 123 | 141 | 121 | 55 | 85 | 1 |
| 109 | 47 | 79 | 106 | 105 | 120 | 127 | 4 |
| 98 | 49 | 141 | 127 | 106 | 82 | 86 | 2 |
| 64 | 114 | 97 | 173 | 151 | 53 | 79 | 2 |
| 139 | 84 | 86 | 104 | 133 | 128 | 86 | 5 |
| 103 | 133 | 88 |  | 129 | 83 |  | 2 |
| 116 | 159 |  |  |  |  |  |  |
| 133 | 5 |  |  |  |  |  |  |
| 31 | 131 |  |  |  |  |  |  |
| 85 | 80 |  |  |  |  |  |  |
| 144 | 112 |  |  |  |  |  |  |
| 83 | 100 |  |  |  |  |  |  |
| 82 | 111 |  |  |  |  |  |  |
| 77 | 66 |  |  |  |  |  |  |
| 70 | 99 |  |  |  |  |  |  |
| 92 | 85 |  |  |  |  |  |  |
| 147 | 113 |  |  |  |  |  |  |
| 98 | 92 |  |  |  |  |  |  |
| 128 | 91 |  |  |  |  |  |  |

**Table EV4:** Inflammatory response of HD mice following chronic injection of hybrid g7-NPs-chol (“2-cycle treatment”, *N* = 4/group). Analytes whose values are significantly different in zQ175DN+chol group compared to control group are in red boxes. Data information: data are expressed as mean±SEM. Statistics: one-way ANOVA with Tuckey post-hoc test (*p<0.05; **p<0.01).

| Analyte | Striatum (vs wt) |  |  |  | Cortex (vs wt) |  |  |  | Cerebellum (vs wt) |  |  |  |
| --- | --- | --- | --- | --- | --- | --- | --- | --- | --- | --- | --- | --- |
|  | zQ175DN |  | zQ175DN+chol |  | zQ175DN |  | zQ175DN+chol |  | zQ175DN |  | zQ175DN+chol |  |
|  | Mean | SEM | Mean | SEM | Mean | SEM | Mean | SEM | Mean | SEM | Mean | SEM |
| IL-1a | 0,722 | 0,056 | 0,667 | 0,000 | 0,778 | 0,064 | 0,556 | 0,064 | 0,895 | 0,053 | 1,053 | 0,086 |
| IL-1b | 0,909 | 0,105 | 0,546 | 0,105 | 0,700 | 0,100 | 0,650 | 0,126 | 1,600 | 0,432 | 1,500 | 0,100 |
| IL-2 | 0,870 | 0,039 | 0,610 | 0,100 | 1,010 | 0,172 | 0,711 | 0,062 | 1,228 | 0,198 | 1,519 | 0,216 |
| IL-3 | N/D |  | N/D |  | N/D |  | N/D |  | N/D |  | N/D |  |
| IL-4 | N/D |  | N/D |  | N/D |  | N/D |  | N/D |  | N/D |  |
| IL-5 | 1,000 | 0,577 | 1,000 | 0,000 | 0,500 | 0,289 | 0,333 | 0,333 | 0,857 | 0,286 | 0,429 | 0,143 |
| IL-6 | 0,135 | 0,037 | 0,077 | 0,000 | 0,750 | 0,144 | 0,375 | 0,125 | 0,364 | 0,149 | 0,364 | 0,000 |
| IL-9 | 0,770 | 0,068 | 0,892 | 0,126 | 0,641 | 0,078 | 0,719 | 0,131 | 0,761 | 0,043 | 0,682 | 0,049 |
| IL-10 | 0,783 | 0,112 | 0,522 | 0,142 | 1,087 | 0,336 | 0,522 | 0,159 | 0,931 | 0,335 | 0,724 | 0,153 |
| IL-12 (p40) | 0,636 | 0,189 | 0,500 | 0,087 | 0,571 | 0,117 | 0,476 | 0,095 | 0,400 | 0,103 | 0,320 | 0,065 |
| IL-12 (p70) | 0,622 | 0,140 | 0,437 | 0,033 | 0,663 | 0,156 | 1,375 | 0,821 | 0,800 | 0,063 | 1,064 | 0,215 |
| IL-13 | 0,625 | 0,052 | 0,627 | 0,103 | 0,896 | 0,176 | 0,701 | 0,033 | 1,452 | 0,131 | 1,305 | 0,223 |
| IL-17 | 0,595 | 0,114 | 0,417 | 0,049 | 0,756 | 0,253 | 0,467 | 0,076 | 0,596 | 0,042 | 0,404 | 0,057 |
| Eotaxin | 0,719 | 0,060 | 0,469 * | 0,031 | 0,722 | 0,106 | 0,639 | 0,070 | 0,878 | 0,138 | 0,781 | 0,040 |
| G-CSF | 0,849 | 0,121 | 0,636 | 0,091 | 0,500 | 0,000 | 0,750 | 0,250 | 0,467 | 0,128 | 0,533 | - |
| GM-CSF | 0,804 | 0,079 | 0,768 | 0,034 | 0,867 | 0,147 | 0,667 | 0,077 | 0,754 | 0,019 | 0,672 | 0,049 |
| ING-gamma | 0,500 | 0,084 | 0,466 | 0,028 | 0,897 | 0,119 | 0,588 | 0,120 | 0,607 | 0,031 | 0,533 | 0,050 |
| KC | 0,594 | 0,094 | 0,375 ** | 0,000 | 0,571 | 0,078 | 0,571 | 0,078 | 0,553 | 0,050 | 0,447 * | 0,079 |
| MCP-1 | 0,068 | 0,005 | 0,093 | 0,017 | 0,899 | 0,232 | 0,930 | 0,250 | 0,843 | 0,126 | 0,825 | 0,127 |
| MIP-1a | 1,047 | 0,114 | 1,043 | 0,117 | 1,167 | 0,271 | 0,892 | 0,113 | 0,713 | 0,116 | 0,629 | 0,117 |
| MIP-1b | 0,737 | 0,048 | 0,618 | 0,142 | 0,909 | 0,235 | 0,212 | 0,030 | 0,518 | 0,080 | 0,304 | 0,042 |
| Rantes | 0,680 | 0,067 | 0,530 | 0,075 | 0,983 | 0,178 | 1,017 | 0,144 | 1,388 | 0,194 | 1,327 | 0,212 |
| TNF-a | 0,610 | 0,028 | 0,695 | 0,198 | 0,725 | 0,180 | 0,975 | 0,170 | 0,559 | 0,093 | 0,509 | 0,081 |

| Analyte | Lung (vs wt) |  |  |  | Liver (vs wt) |  |  |  | Kidney (vs wt) |  |  |  |
| --- | --- | --- | --- | --- | --- | --- | --- | --- | --- | --- | --- | --- |
|  | zQ175DN |  | zQ175DN+chol |  | zQ175DN |  | zQ175DN+chol |  | zQ175DN |  | zQ175DN+chol |  |
|  | Mean | SEM | Mean | SEM | Mean | SEM | Mean | SEM | Mean | SEM | Mean | SEM |
| IL-1a | N/D |  | N/D |  | 0,680 | 0,214 | 0,760 | 0,090 | 0,874 | 0,036 | 1,207 | 0,174 |
| IL-1b | 0,667 | 0,000 | 0,667 | 0,000 | 1,200 | 0,300 | 0,900 | 0,000 | 1,167 | 0,167 | 1,000 | 0,000 |
| IL-2 | 0,692 | 0,000 | 1,578 | 0,451 | 1,447 | 0,417 | 1,303 | 0,224 | 0,919 | 0,193 | 0,730 | - |
| IL-3 | N/D |  | N/D |  | N/D |  | N/D |  | N/D |  | N/D |  |
| IL-4 | N/D |  | N/D |  | N/D |  | N/D |  | N/D |  | N/D |  |
| IL-5 | 0,762 | 0,098 | 0,434 | 0,067 | 1,371 | 0,851 | 1,777 | 0,390 | 0,671 | 0,095 | 0,710 | 0,065 |
| IL-6 | 0,718 | 0,271 | 1,231 | 0,774 | 0,913 | 0,126 | 0,543 | 0,174 | 0,752 | 0,068 | 1,282 | 0,600 |
| IL-9 | 1,415 | 0,434 | 1,189 | 1,083 | 0,658 | 0,045 | 0,518 | 0,020 | 0,410 | 0,035 | 0,519 | - |
| IL-10 | 1,000 | 0,406 | 0,878 | 0,378 | 0,734 | 0,008 | 0,677 | 0,000 | 1,104 | 0,203 | 0,816 | 0,190 |
| IL-12 (p40) | 0,998 | 0,000 | 0,998 | 0,003 | 0,992 | 0,001 | 0,986 | 0,015 | 0,996 | 0,005 | 0,993 | 0,006 |
| IL-12 (p70) | 0,858 | 0,024 | 1,135 | 0,256 | 1,028 | 0,054 | 0,942 | 0,111 | 0,818 | 0,091 | 0,679 | 0,032 |
| IL-13 | 0,998 | 0,141 | 0,977 | 0,028 | 1,023 | 0,064 | 0,524 | 0,213 | 0,637 | 0,565 | 1,290 | 0,113 |
| IL-17 | 0,889 | 0,222 | 0,833 | 0,096 | 0,583 | 0,083 | 1,000 | 0,250 | 0,667 | 0,000 | 0,778 | 0,111 |
| Eotaxin | 0,728 | 0,264 | 0,779 | 0,222 | 0,847 | 0,086 | 0,824 | 0,272 | 0,810 | 0,160 | 0,625 | 0,089 |
| G-CSF | 0,937 | 0,131 | 1,104 | 0,127 | 0,741 | 0,157 | 0,608 | 0,118 | 2,470 | 1,700 | 0,752 | 0,127 |
| GM-CSF | 0,815 | 0,074 | 0,926 | 0,134 | 1,193 | 0,153 | 1,105 | 0,158 | 1,279 | 0,154 | 0,930 | 0,000 |
| ING-gamma | 0,893 | 0,114 | 1,276 | 0,155 | 0,802 | 0,159 | 0,649 | 0,395 | 0,785 | 0,175 | 0,734 | 0,142 |
| KC | 0,639 | 0,083 | 0,635 | 0,035 | 0,654 | 0,116 | 0,438 * | 0,132 | 0,646 | 0,098 | 0,813 | 0,018 |
| MCP-1 | 1,153 | 0,133 | 1,063 | 0,509 | 0,905 | 0,171 | 0,947 | 0,213 | 0,924 | 0,104 | 0,930 | 0,174 |
| MIP-1a | N/D |  | N/D |  | N/D |  | N/D |  | N/D |  | N/D |  |
| MIP-1b | 0,638 | 0,185 | 0,185 | 0,021 | 0,816 | 0,037 | 0,521 | 0,160 | 1,024 | 0,397 | 1,250 | 0,541 |
| Rantes | 0,743 | 0,257 | 1,004 | 0,075 | 0,944 | 0,100 | 0,738 | 0,310 | 1,141 | 0,131 | 1,227 | 0,272 |
| TNF-a | 0,881 | 0,346 | 1,220 | 0,409 | 1,191 | 0,409 | 0,454 | 0,331 | 0,917 | 0,027 | 0,684 | - |

| Analyte | Spleen (vs wt) |  |  |  | Heart (vs wt) |  |  |  | Plasma (vs wt) |  |  |  |
| --- | --- | --- | --- | --- | --- | --- | --- | --- | --- | --- | --- | --- |
|  | zQ175DN |  | zQ175DN+chol |  | zQ175DN |  | zQ175DN+chol |  | zQ175DN |  | zQ175DN+chol |  |
|  | Mean | SEM | Mean | SEM | Mean | SEM | Mean | SEM | Mean | SEM | Mean | SEM |
| IL-1a | 0,772 | 0,140 | 0,579 | 0,101 | 0,750 | 0,205 | 1,071 | 0,327 | 1,250 | 0,433 | 1,125 | 0,389 |
| IL-1b | 1,208 | 0,386 | 1,198 | 0,361 | 0,735 | 0,062 | 0,525 | 0,149 | 1,387 | - | 0,465 | 0,070 |
| IL-2 | 0,750 | 0,076 | 0,804 | 0,103 | 0,928 | 0,226 | 0,978 | 0,125 | 0,546 | - | 1,455 | - |
| IL-3 | N/D |  | N/D |  | N/D |  | N/D |  | N/D |  | N/D |  |
| IL-4 | N/D |  | N/D |  | N/D |  | N/D |  | N/D |  | N/D |  |
| IL-5 | N/D |  | N/D |  | 0,750 | 0,000 | 1,250 | 0,500 | 0,476 | 0,095 | 0,476 | 0,286 |
| IL-6 | 0,144 | 0,011 | 0,167 | 0,056 | 1,350 | 0,150 | 0,900 | 0,000 | 0,800 | 0,267 | 0,978 | 0,470 |
| IL-9 | 1,250 | 0,250 | 0,889 | 0,111 | 0,667 | 0,000 | 0,945 | 0,278 | 0,750 | 0,250 | 0,500 | - |
| IL-10 | N/D |  | N/D |  | 1,115 | 0,366 | 0,847 | 0,191 | 0,667 | 0,273 | 0,404 | - |
| IL-12 (p40) | N/D |  | N/D |  | N/D |  | N/D |  | N/D |  | N/D |  |
| IL-12 (p70) | 1,520 | 0,843 | 0,423 | 0,141 | 1,386 | 0,486 | 0,477 | 0,000 | 2,291 | 1,264 | 1,156 | 0,345 |
| IL-13 | 1,018 | 0,296 | 0,718 | 0,131 | 1,091 | 0,105 | 0,909 | 0,065 | 0,788 | 0,290 | 0,575 | 0,084 |
| IL-17 | 1,905 | 0,780 | 0,643 | 0,270 | 1,071 | 0,643 | 0,643 | 0,000 | 0,333 | - | 0,556 | 0,222 |
| Eotaxin | N/D |  | N/D |  | N/D |  | N/D |  | N/D |  | N/D |  |
| G-CSF | 0,500 | 0,000 | 0,688 | 0,120 | 3,000 | 2,000 | 1,000 | 0,000 | 0,366 | - | 1,162 | 0,791 |
| GM-CSF | 1,282 | 0,303 | 1,016 | 0,071 | 0,989 | 0,033 | 0,935 | 0,042 | 1,062 | 0,122 | 0,943 | 0,035 |
| ING-gamma | 0,866 | 0,082 | 0,698 | 0,119 | 3,118 | - | 1,482 | 0,581 | 0,779 | 0,095 | 0,945 | 0,025 |
| KC | 0,885 | 0,128 | 0,695 | 0,320 | 0,826 | 0,242 | 0,776 | 0,064 | 0,558 | 0,315 | 0,781 | 0,439 |
| MCP-1 | 0,921 | 0,020 | 0,927 | 0,091 | 0,919 | 0,029 | 0,740 | 0,048 | 0,619 | 0,095 | 0,582 | 0,018 |
| MIP-1a | 1,000 | 0,250 | 0,656 | 0,054 | N/D |  | N/D |  | 0,667 | 0,381 | 0,429 | 0,143 |
| MIP-1b | 0,956 | 0,163 | 1,014 | 0,330 | 0,801 | 0,082 | 0,852 | 0,048 | 2,652 | 0,967 | 1,605 | 0,310 |
| Rantes | 2,097 | 0,768 | 0,247 | - | 0,741 | 0,158 | 0,685 | 0,067 | 1,667 | 0,500 | 2,500 | 0,536 |
| TNF-a | 1,087 | 0,513 | 1,017 | 0,402 | 0,845 | 0,027 | 0,938 | 0,177 | 1,229 | 0,263 | 0,836 | 0,296 |

**Table EV5:**
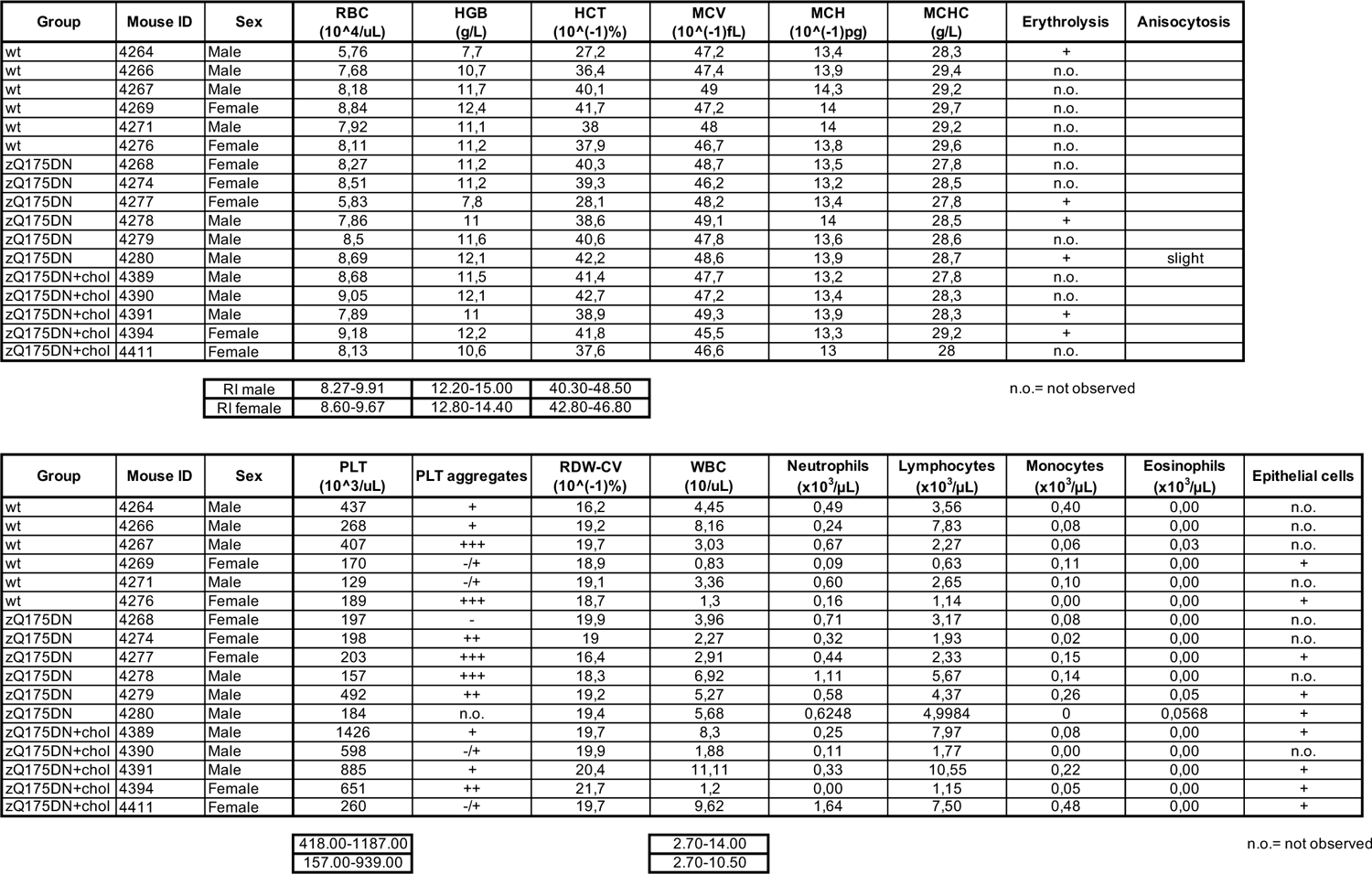
Complete blood count of wt and HD mice following systemic and chronic injection of hybrid g7-NPs-chol (“2-cycle treatment”, *N* = 5-6/group). R.I.= reference intervals. Statistics: Kruskal-Wallis followed by Mann-Whitney test.

**Table EV6:**
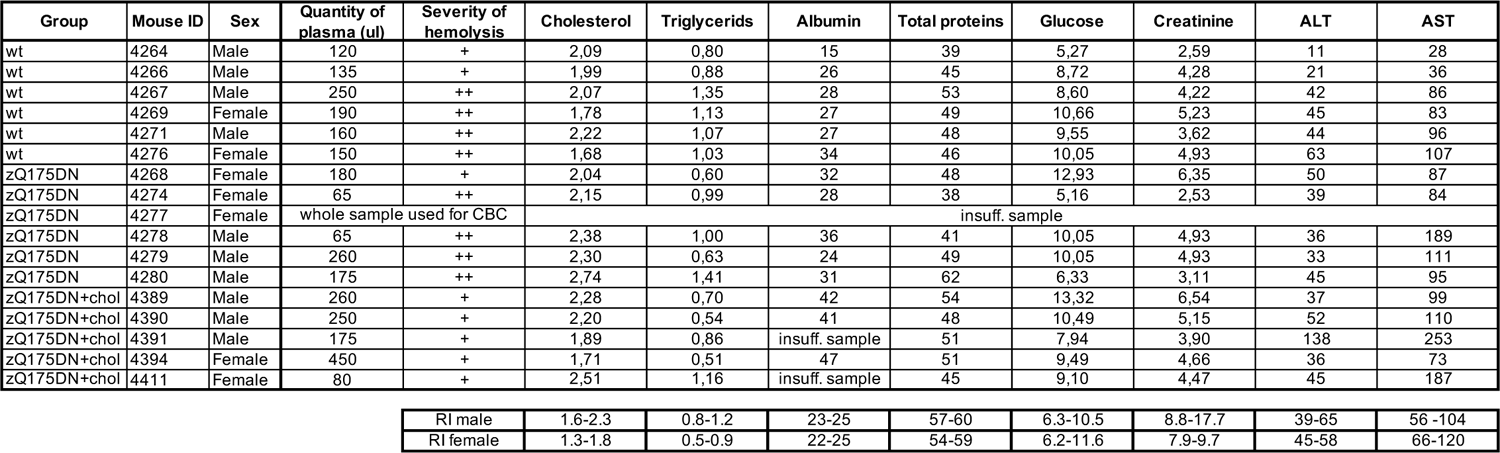
Blood chemistry of wt and HD mice following systemic and chronic injection of hybrid g7-NPs-chol (“2-cycle treatment”, *N* = 5-6/group). CBC = complete blood count; R.I.= reference intervals. Statistics: Kruskal-Wallis followed by Mann-Whitney test.

| Group | Mouse ID | Sex | Quantity of plasma (ul) | Severity of hemolysis | Cholesterol | Triglycerids | Albumin | Total proteins | Glucose | Creatinine | ALT | AST |
| --- | --- | --- | --- | --- | --- | --- | --- | --- | --- | --- | --- | --- |
| wt | 4264 | Male | 120 | + | 2,09 | 0,80 | 15 | 39 | 5,27 | 2,59 | 11 | 28 |
| wt | 4266 | Male | 135 | + | 1,99 | 0,88 | 26 | 45 | 8,72 | 4,28 | 21 | 36 |
| wt | 4267 | Male | 250 | ++ | 2,07 | 1,35 | 28 | 53 | 8,60 | 4,22 | 42 | 86 |
| wt | 4269 | Female | 190 | ++ | 1,78 | 1,13 | 27 | 49 | 10,66 | 5,23 | 45 | 83 |
| wt | 4271 | Male | 160 | ++ | 2,22 | 1,07 | 27 | 48 | 9,55 | 3,62 | 44 | 96 |
| wt | 4276 | Female | 150 | ++ | 1,68 | 1,03 | 34 | 46 | 10,05 | 4,93 | 63 | 107 |
| zQ175DN | 4268 | Female | 180 | + | 2,04 | 0,60 | 32 | 48 | 12,93 | 6,35 | 50 | 87 |
| zQ175DN | 4274 | Female | 65 | ++ | 2,15 | 0,99 | 28 | 38 | 5,16 | 2,53 | 39 | 84 |
| zQ175DN | 4277 | Female | whole sample used for CBC |  | insuff. sample |  |  |  |  |  |  |  |
| zQ175DN | 4278 | Male | 65 | ++ | 2,38 | 1,00 | 36 | 41 | 10,05 | 4,93 | 36 | 189 |
| zQ175DN | 4279 | Male | 260 | ++ | 2,30 | 0,63 | 24 | 49 | 10,05 | 4,93 | 33 | 111 |
| zQ175DN | 4280 | Male | 175 | ++ | 2,74 | 1,41 | 31 | 62 | 6,33 | 3,11 | 45 | 95 |
| zQ175DN+chol | 4389 | Male | 260 | + | 2,28 | 0,70 | 42 | 54 | 13,32 | 6,54 | 37 | 99 |
| zQ175DN+chol | 4390 | Male | 250 | + | 2,20 | 0,54 | 41 | 48 | 10,49 | 5,15 | 52 | 110 |
| zQ175DN+chol | 4391 | Male | 175 | + | 1,89 | 0,86 | insuff. sample | 51 | 7,94 | 3,90 | 138 | 253 |
| zQ175DN+chol | 4394 | Female | 450 | + | 1,71 | 0,51 | 47 | 51 | 9,49 | 4,66 | 36 | 73 |
| zQ175DN+chol | 4411 | Female | 80 | + | 2,51 | 1,16 | insuff. sample | 45 | 9,10 | 4,47 | 45 | 187 |

|  |  |  |  |  |  |  |  |  |
| --- | --- | --- | --- | --- | --- | --- | --- | --- |
| RI male | 1.6-2.3 | 0.8-1.2 | 23-25 | 57-60 | 6.3-10.5 | 8.8-17.7 | 39-65 | 56 -104 |
| RI female | 1.3-1.8 | 0.5-0.9 | 22-25 | 54-59 | 6.2-11.6 | 7.9-9.7 | 45-58 | 66-120 |

**Table EV7:** Body weight and weight of organs in grams of wt and HD mice following systemic and chronic injection of hybrid g7-NPs-chol (“2-cycle treatment”, *N* = 5-6/group). Statistics: Kruskal-Wallis followed by Mann-Whitney test.

| Group | Mouse ID | Sex | Weight (g) |  |  |  |  |  |  |  |
| --- | --- | --- | --- | --- | --- | --- | --- | --- | --- | --- |
|  |  |  | Body | Liver | Spleen | Left kidney | Right kidney | Heart | Left testis | Right testis |
| wt | 4264 | Male | 28,7 | 1,680 | 0,107 | 0,348 | 0,773 | 0,215 | 0,083 | 0,090 |
| wt | 4266 | Male | 31,1 | 1,603 | 0,087 | 0,207 | 0,190 | 0,159 | 0,090 | 0,086 |
| wt | 4267 | Male | 32,2 | 1,715 | 0,096 | 0,226 | 0,220 | 0,205 | 0,093 | 0,087 |
| wt | 4269 | Female | 27,7 | 1,554 | 0,120 | 0,182 | 0,176 | 0,153 | / | / |
| wt | 4271 | Male | 36,4 | 1,910 | 0,092 | 0,238 | 0,235 | 0,243 | 0,102 | 0,113 |
| wt | 4276 | Female | 26,7 | 1,455 | 0,112 | 0,141 | 0,161 | 0,187 | / | / |
| zQ175DN | 4268 | Female | 22,3 | 1,087 | 0,117 | 0,126 | 0,140 | 0,136 | / | / |
| zQ175DN | 4274 | Female | 22,2 | 1,129 | 0,096 | 0,133 | 0,132 | 0,144 | / | / |
| zQ175DN | 4277 | Female | 22,8 | 1,168 | 0,113 | 0,141 | 0,142 | 0,130 | / | / |
| zQ175DN | 4278 | Male | 26,7 | 1,417 | 0,072 | 0,167 | 0,168 | 0,139 | 0,092 | 0,092 |
| zQ175DN | 4279 | Male | 29,3 | 1,542 | 0,083 | 0,171 | 0,182 | 0,144 | 0,100 | 0,108 |
| zQ175DN | 4280 | Male | 32,0 | 1,637 | 0,080 | 0,197 | 0,208 | 0,191 | 0,096 | 0,091 |
| zQ175DN+chol | 4389 | Male | 26,3 | 1,375 | 0,063 | 0,144 | 0,150 | 0,141 | 0,088 | 0,111 |
| zQ175DN+chol | 4390 | Male | 25,4 | 1,415 | 0,094 | 0,162 | 0,170 | 0,149 | 0,100 | 0,091 |
| zQ175DN+chol | 4391 | Male | 25,4 | 1,021 | 0,095 | 0,205 | 0,221 | 0,165 | 0,090 | 0,090 |
| zQ175DN+chol | 4394 | Female | 20,8 | 1,180 | 0,116 | 0,127 | 0,127 | 0,124 | / | / |
| zQ175DN+chol | 4411 | Female | 20,8 | 1,158 | 0,096 | 0,129 | 0,129 | 0,116 | / | / |

**Table EV8:**
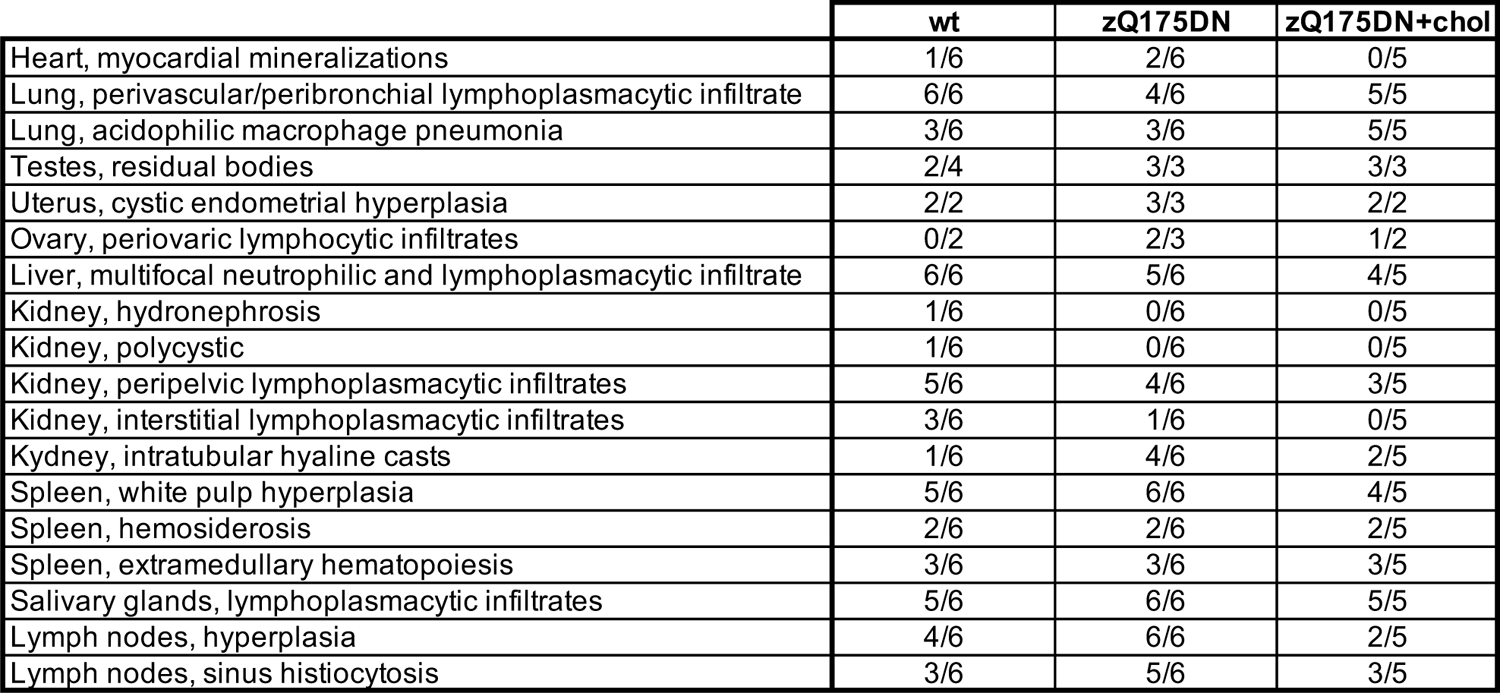
Summary of gross and histological lesions of wt and HD mice following systemic and chronic injection of hybrid g7-NPs-chol (“2-cycle treatment”, *N* = 5-6/group).

**Table EV9:** Foreign body granuloma analysis of HD mice following systemic and chronic injection of hybrid g7-NPs-chol for all the treatments.

| Type of NPs | N° of IP injections | Weeks after the last IP | N° of nodules | Concretions diameter |
| --- | --- | --- | --- | --- |
| g7-hybrid-NPs | 10 | 20 | 1 (not in all mice) | 1-2 mm |
| g7-hybrid-NPs | 10 | 40 | 1-2 (not in all mice) | 1-2 mm |
| g7-hybrid-NPs | 10 + 10 (2 cycles) | 20 | 5-6 (in all mice) | 2-3 mm |

## REFERENCES

1. Martín MG, Pfrieger F & Dotti CG (2014) Cholesterol in brain disease: sometimes determinant and frequently implicated. EMBO Rep 15: 1036–1052

2. Saudou F & Humbert S (2016) The Biology of Huntingtin. Neuron 89: 910–926

3. Zuccato C, Valenza M & Cattaneo E (2010) Molecular Mechanisms and Potential Therapeutical Targets in Huntington’s Disease. Physiol Rev 90: 905–981

4. Rüb U, Seidel K, Heinsen H, Vonsattel JP, den Dunnen WF & Korf HW (2016) Huntington’s disease (HD): the neuropathology of a multisystem neurodegenerative disorder of the human brain. Brain Pathology 26: 726–740

5. Valenza M, Leoni V, Tarditi A, Mariotti C, Björkhem I, Di Donato S & Cattaneo E (2007a) Progressive dysfunction of the cholesterol biosynthesis pathway in the R6/2 mouse model of Huntington’s disease. Neurobiol Dis 28: 133–142

6. Valenza M, Carroll JB, Leoni V, Bertram LN, Björkhem I, Singaraja RR, Di Donato S, Lutjohann D, Hayden MR & Cattaneo E (2007b) Cholesterol biosynthesis pathway is disturbed in YAC128 mice and is modulated by huntingtin mutation. Hum Mol Genet 16: 2187–2198

7. Kacher R, Lamazière A, Heck N, Kappes V, Mounier C, Despres G, Dembitskaya Y, Perrin E, Christaller W, Sasidharan Nair S, et al. (2019) CYP46A1 gene therapy deciphers the role of brain cholesterol metabolism in Huntington’s disease. Brain 142: 2432–2450

8. Shankaran M, Di Paolo E, Leoni V, Caccia C, Ferrari Bardile C, Mohammed H, Di Donato S, Kwak S, Marchionini D, Turner S, et al. (2017) Early and brain region-specific decrease of de novo cholesterol biosynthesis in Huntington’s disease: A cross-validation study in Q175 knock-in mice. Neurobiol Dis 98: 66–76

9. Jurevics H & Morell P (1995) Cholesterol for synthesis of myelin is made locally, not imported into brain. Journal of Neurochemistry 64: 895–901

10. Björkhem I, Meaney S & Fogelman AM (2004) Brain Cholesterol: Long Secret Life behind a Barrier. Arterioscler Thromb Vasc Biol 24: 806–815

11. Li D, Zhang J & Liu Q (2022) Brain cell type-specific cholesterol metabolism and implications for learning and memory. Trends Neurosci 45: 401–414

12. Leoni V, Mariotti C, Tabrizi SJ, Valenza M, Wild EJ, Henley SMD, Hobbs NZ, Mandelli ML, Grisoli M, Björkhem I, et al. (2008) Plasma 24S-hydroxycholesterol and caudate MRI in pre-manifest and early Huntington’s disease. Brain 131: 2851–2859

13. Leoni V, Mariotti C, Nanetti L, Salvatore E, Squitieri F, Bentivoglio AR, Bandettini del Poggio M, Piacentini S, Monza D, Valenza M, et al. (2011) Whole body cholesterol metabolism is impaired in Huntington’s disease. Neurosci Lett 494: 245–249

14. Leoni V, Long JD, Mills JA, Di Donato S & Paulsen JS (2013) Plasma 24S-hydroxycholesterol correlation with markers of Huntington disease progression. Neurobiol Dis 55: 37–43

15. Tosi G, Costantino L, Rivasi F, Ruozi B, Leo E, Vergoni A V., Tacchi R, Bertolini A, Vandelli MA & Forni F (2007) Targeting the central nervous system: In vivo experiments with peptide-derivatized nanoparticles loaded with Loperamide and Rhodamine-123. Journal of Controlled Release 122: 1–9

16. Valenza M, Chen JY, Di Paolo E, Ruozi B, Belletti D, Ferrari Bardile C, Leoni V, Caccia C, Brilli E, Di Donato S, et al. (2015a) Cholesterol-loaded nanoparticles ameliorate synaptic and cognitive function in H untington’s disease mice. EMBO Mol Med 7: 1547–1564

17. Birolini G, Valenza M, Di Paolo E, Vezzoli E, Talpo F, Maniezzi C, Caccia C, Leoni V, Taroni F, Bocchi VD, et al. (2020) Striatal infusion of cholesterol promotes dose-dependent behavioral benefits and exerts disease-modifying effects in Huntington’s disease mice. EMBO Mol Med 12:e12519.

18. Birolini G, Verlengia G, Talpo F, Maniezzi C, Zentilin L, Giacca M, Conforti P, Cordiglieri C, Caccia C, Leoni V, et al. (2021a) SREBP2 gene therapy targeting striatal astrocytes ameliorates Huntington’s disease phenotypes. Brain 144: 3175–3190

19. Belletti D, Grabrucker AM, Pederzoli F, Menerath I, Vandelli MA, Tosi G, Duskey TJ, Forni F & Ruozi B (2018) Hybrid nanoparticles as a new technological approach to enhance the delivery of cholesterol into the brain. Int J Pharm 543: 300–310

20. Birolini G, Valenza M, Ottonelli I, Passoni A, Favagrossa M, Duskey JT, Bombaci M, Vandelli MA, Colombo L, Bagnati R, et al. (2021b) Insights into kinetics, release, and behavioral effects of brain-targeted hybrid nanoparticles for cholesterol delivery in Huntington’s disease. Journal of Controlled Release 330: 587–598

21. Menalled LB, Kudwa AE, Miller S, Fitzpatrick J, Watson-Johnson J, Keating N, Ruiz M, Mushlin R, Alosio W, McConnell K, et al. (2012) Comprehensive Behavioral and Molecular Characterization of a New Knock-In Mouse Model of Huntington’s Disease: ZQ175. PLoS One 7: e49838

22. Southwell AL, Smith-Dijak A, Kay C, Sepers M, Villanueva EB, Parsons MP, Xie Y, Anderson L, Felczak B, Waltl S, et al. (2016) An enhanced Q175 knock-in mouse model of Huntington disease with higher mutant huntingtin levels and accelerated disease phenotypes. Hum Mol Genet 25: 3654–3675

23. Tosi G, Vilella A, Chhabra R, Schmeisser MJ, Boeckers TM, Ruozi B, Vandelli MA, Forni F, Zoli M & Grabrucker AM (2014) Insight on the fate of CNS-targeted nanoparticles. Part II: Intercellular neuronal cell-to-cell transport. Journal of Controlled Release 177: 96–107

24. Cepeda C, Wu N, André VM, Cummings DM & Levine MS (2007) The Corticostriatal Pathway in Huntington’s Disease. Prog Neurbiol 81:253–271

25. Milnerwood AJ & Raymond LA (2010) Early synaptic pathophysiology in neurodegeneration: Insights from Huntington’s disease. Trends Neurosci 33: 513–523

26. Heikkinen T, Lehtimäki K, Vartiainen N, Puoliväli J, Hendricks SJ, Glaser JR, Bradaia A, Wadel K, Touller C, Kontkanen O, et al. (2012) Characterization of Neurophysiological and Behavioral Changes, MRI Brain Volumetry and 1H MRS in zQ175 Knock-In Mouse Model of Huntington’s Disease. PLoS One 7: e50717

27. Indersmitten T, Tran CH, Cepeda C, Levine MS (2015) Altered excitatory and inhibitory inputs to striatal medium-sized spiny neurons and cortical pyramidal neurons in the Q175 mouse model of Huntington’s disease. Journal of Neurophysiology 113: 2953–2966

28. Vezzoli E, Caron I, Talpo F, Besusso D, Conforti P, Battaglia E, Sogne E, Falqui A, Petricca L, Verani M, et al. (2019) Inhibiting pathologically active ADAM10 rescues synaptic and cognitive decline in Huntington’s disease. Journal of Clinical Investigation 129: 2390–2403

29. Passoni A, Favagrossa M, Colombo L, Bagnati R, Gobbi M, Diomede L, Birolini G, Di Paolo E, Valenza M, Cattaneo E, et al. (2020) Efficacy of Cholesterol Nose-to-Brain Delivery for Brain Targeting in Huntington’s Disease. ACS Chem Neurosci 11: 367–372

30. Valenza M, Marullo M, Di Paolo E, Cesana E, Zuccato C, Biella G & Cattaneo E (2015b) Disruption of astrocyte-neuron cholesterol cross talk affects neuronal function in Huntington’s disease. Cell Death Differ 22: 690–702

31. Boussicault L, Alves S, Lamazière A, Planques A, Heck N, Moumné L, Despres G, Bolte S, Hu A, Pagès C, et al. (2016) CYP46A1, the rate-limiting enzyme for cholesterol degradation, is neuroprotective in Huntington’s disease. Brain 139: 953–970

32. Valenza M, Leoni V, Karasinska JM, Petricca L, Fan J, Carroll J, Pouladi MA, Fossale E, Nguyen HP, Riess O, et al. (2010) Cholesterol defect is marked across multiple rodent models of Huntington’s disease and is manifest in astrocytes. Journal of Neuroscience 30: 10844–10850

33. Kordasiewicz HB, Stanek LM, Wancewicz E V., Mazur C, McAlonis MM, Pytel KA, Artates JW, Weiss A, Cheng SH, Shihabuddin LS, et al. (2012) Sustained Therapeutic Reversal of Huntington’s Disease by Transient Repression of Huntingtin Synthesis. Neuron 74: 1031–1044

34. Zeitler B, Froelich S, Marlen K, Shivak DA, Yu Q, Li D, Pearl JR, Miller JC, Zhang L, Paschon DE, et al. (2019) Allele-selective transcriptional repression of mutant HTT for the treatment of Huntington’s disease. Nat Med 25: 1131–1142

35. Pikuleva IA & Cartier N (2021) Cholesterol Hydroxylating Cytochrome P450 46A1: From Mechanisms of Action to Clinical Applications. Front Aging Neurosci 13: 696778

36. Marchionini DM, Liu J-P, Ambesi-Impiombato A, Kerker K, Cirillo K, Bansal M, Mushlin R, Brunner D, Ramboz S, Kwan M, et al (2022) Benefits of global mutant huntingtin lowering diminish over time in a Huntington’s disease mouse model. JCI Insight 7: e161769

37. Rao AK, Gordon AM & Marder KS (2011) Coordination of fingertip forces during precision grip in premanifest Huntington’s disease. Movement Disorders 26: 862–869

38. Paulsen JS, Nance M, Kim JI, Carlozzi NE, Panegyres PK, Erwin C, Goh A, McCusker E & Williams JK (2013) A review of quality of life after predictive testing for and earlier identification of neurodegenerative diseases. Prog Neurobiol 110: 2–28

39. Ramos ARS & Garrett C (2017) Huntington’s Disease: Premotor Phase. Neurodegener Dis 17: 313– 322

40. Darvas M & Palmiter RD (2009) Restriction of dopamine signaling to the dorsolateral striatum is sufficient for many cognitive behaviors Proc Natl Acad Sci U S A 106: 14664–14669

41. Poulter SL, Kosaki Y, Sanderson DJ & McGregor A (2020) Spontaneous object-location memory based on environmental geometry is impaired by both hippocampal and dorsolateral striatal lesions. Brain Neurosci Adv 4: 239821282097259

42. Krabbe S, Gründemann J & Lüthi A (2018) Amygdala Inhibitory Circuits Regulate Associative Fear Conditioning. Biol Psychiatry 83: 800–809

43. Goodroe SC, Starnes J & Brown TI (2018) The Complex Nature of Hippocampal-Striatal Interactions in Spatial Navigation. Front Hum Neurosci 12: 250

44. Abildayeva K, Jansen PJ, Hirsch-Reinshagen V, Bloks VW, Bakker AHF, Ramaekers FCS, De Vente J, Groen AK, Wellington CL, Kuipers F, et al. (2006) 24(S)-hydroxycholesterol participates in a liver X receptor-controlled pathway in astrocytes that regulates apolipoprotein E-mediated cholesterol efflux. Journal of Biological Chemistry 281: 12799–12808

45. Ramot Y, Ben-Eliahu S, Kagan L, Ezov N & Nyska A (2009) Subcutaneous and Intraperitoneal Lipogranulomas Following Subcutaneous Injection of Olive Oil in Sprague-Dawley Rats. Toxicol Pathol 37: 882–886

46. Alsina-Sanchis E, Mülfarth R, Moll I, Mogler C, Rodriguez-Vita J & Fischer A (2021) Intraperitoneal oil application causes local inflammation with depletion of resident peritoneal macrophages. Molecular Cancer Research 19: 288–300

